# Longest Order Conserved Exemplar Subsequences

**DOI:** 10.1101/2020.12.15.422841

**Authors:** Shu Zhang, Lianrong Pu, Runmin Yang, Luli Wang, Daming Zhu, Haitao Jiang

## Abstract

We propose a new problem whose input data are two linear genomes together with two indexed gene subsequences of them, which asks to find a longest common exemplar subsequence of the two given genomes with a subsequence identical to the given indexed gene subsequences. We present an algorithm for this problem such that the algorithm is allowed to take diminishing time and space to solve the problem by setting the indexed genes with an incremental number. Although an incremental number of indexed genes were selected, the algorithm was verified definite to reach a solution whose length insistently comes very close to a real longest common exemplar subsequence of the two given genomes.

Aiming at 23 human/gorilla chromosome pairs, the algorithm was examined for use in questing for longest common exemplar subsequences whose basic units are annotated genes as well as pseudo genes, namely consecutive DNA subsequences. By contrasting the pseudo gene common exemplar subsequences the algorithm had reached for the human chromosomes 7 and 16 and their gorilla homologues with those annotated genes in the human and gorilla chromosomes, we found more than 1 000 and 500 pseudo genes in the human chromosomes 7 and 16 that occur in the same order as they are in the gorilla chromosomes 7 and 16 and, do not overlap with any annotated gene.

**Author summary:** There is a benefit of the algorithm: It can reach a long enough common exemplar subsequence of two linear genomes in as fast a speed as one requires even if the given genomes would be equipped with too many duplicated genes, which can be done by setting incremental number of indexed genes. We developed a Java software based on the algorithm, that has been available for download on https://github.com/ShuZhang-sdu/LCES.

Only in need to set the indexed gene sequences as null, was it verified successful for our algorithm to obtain the longest common exemplar subsequences of the annotated gene summary pairs extracted from 23 human/gorilla chromosome pairs.

In convenience for researchers to find new motifs or conserved genes, we devoted for the algorithm to quest pseudo gene (i.e. consecutive DNA subsequences) summary pairs of the 23 human/gorilla chromosome pairs for solutions. There are 20 pseudo gene summary pairs whose longest common exemplar subsequences have been found by the algorithm with null indexed gene sequences. The other 3 pseudo gene summary pairs were verified solvable for the algorithm to reach their longest common exemplar subsequences that have to admit subsequences identical to given indexed gene subsequences. There were informed to exist 2 353 and 1 148 pseudo genes in the gorilla chromosome 7 and 16 that occur in the same order as they are in the human chromosome 7 and 16 and, do not overlap with any annotated gene. These pseudo genes should be significant for annotating the human or gorilla genome.

## 1 Introduction

Conserved gene based molecular mechanism analysis happens basically in kinds of bioresearch such as breeding soybean seeds [1] as well as developing drugs or vaccines to beat back corona viruses [2] [3] [4]. Finding conserved gene subsequences in genomes has been an attractive topic in biomedicine, bioinformatics and computer science all the time. A consecutive DNA subsequence is not necessarily an annotated gene if it is conserved. Thus a sequence of conserved consecutive DNA subsequences in a genome sounds to have a broader range of applications such as for people to find motifs in it or illuminate bio-functions of it. An order conserved sequence of conserved consecutive DNA subsequences or genes can be found by solving a combinatorial problem in terms of genome comparison [5] [6] [7].

The exemplar breakpoint distance problem (abbr. EBD) proposed by Sankoff is a pioneering problem for this purpose [5]. Some heuristic algorithms using branch and bound [5] or divide and conquer [8] can be found effective for this problem. This problem is NP-Hard [9] and there are extensive approaches on the computational complexity of this problem [10] [11] [12]. Some more algorithmic progresses for special versions of this problem can be found in the literatures [13] [14]. The decision version of EBD, the zero exemplar breakpoint distance problem (abbr. ZEBD), remains NP-Hard [11] and admits an *O*(*n*^2^1.86^*n*^) time algorithm for the two given genomes both to allow at most 2 identical genes where *n* is the number of genes in the given genomes [15]. Some special versions of ZEBD admit polynomial time algorithms [11] [15] [16].

In the sense of order conserved, a long enough common exemplar subsequence of two or more linear genomes should be better qualified for conserved. The exemplar longest common subsequence problem (abbr. ELCS) was proposed by Bonizzoni *et al.* [6] to quest for a longest common subsequence of two or more linear genomes that contains a gene of each mandatory gene family. This problem is NP-Hard since ZEBD is a special version of it and, admits a polynomial time algorithm for the special version whose input is of two linear genomes with at most three genes of each mandatory family [6]. This problem admits fixed-parameter algorithms for instances of two linear genomes [6].

The repetition-free longest common subsequence problem (abbr. RFLCS) proposed by Adi *et al.* [7] is in fact a special version of ELCS. This problem is NP-Hard since ZEBD remains a special version of it. On the positive side, this problem admits an approximation algorithm with performance ratio equal to the maximum number over all those each of which is that one no larger than the other of two numbers of identical genes of the same gene family that respectively occur in the two genomes [7]. A dynamic programming algorithm was proposed for this problem with a time complexity *O*(*mns*4^*s*^) where *s* is the *span*, namely the maximum distance of identical genes in one of the given genomes and, *m* and *n* are the lengths of the two given genomes [17].

An integer linear programming was proposed by Adi *et al.* [7] for RFLCS. Ferreira and Tjandraatmadja [18] proposed a heuristic based branch-and-cut algorithm for the IP described RFLCS, which was verified efficient for those medium size instances. In terms of parameterized algorithms, Blin *et al.* [19] showed that RFLCS rejects any polynomial kernel but admits a randomized FPT algorithm of *O*(2^*k*^*kmn*) time and *O*(*kmn*) space where *k* is the size of an optimal RFLCS solution and *m* and *n* are the lengths of the two given genomes. More meta-heuristic progresses for RFLCS were proposed and experimentally verified efficient [20] [21] [22].

Although so many algorithmic progresses have been proposed with capacity to find conserved gene subsequences, rare of them was based to arrive at a software that could support the use in finding a sequence of conserved subsequences of real genomes. This implies it fundamentally necessary to explore combinatorial problem models that are available for people to design effective algorithms for finding long enough sequences of conserved genes or consecutive DNA subsequences in genomes. Moreover, there exists no approach in terms of finding an order conserved sequence of conserved consecutive DNA subsequences in genomes from genomic data.

In this paper, in motivation of pursuing a longest common exemplar subsequence with genes that have been fixed conserved, we propose a new problem called *longest common exemplar subsequence with indexed genes* (abbr. LCES-IG). The problem with two input genomes is given by two linear genomes *A* = *A*[1] ... *A*[*m*] and *B* = *B*[1] ... *B*[*n*] and, two indexed gene subsequences *X* = *A*[*x*_1_] ... *A*[*x*_*q*_] and *Y* = *B*[*y*_1_] ... *B*[*y*_*q*_] of *A* and *B* where *X* is identical to *Y*, asks to find a longest common exemplar subsequence of *A* and *B* such that it has a subsequence identical to *X* as well as *Y*.

We present a dynamic programming algorithm for LCES-IG such that, if *A* (resp. *B*) in exclusion of those identical to genes in *X* (resp. *Y*) admits a span *s*(*A*, *X*) (resp. *s*(*B*, *Y*)), then the time complexity and the space complexity of the algorithm are 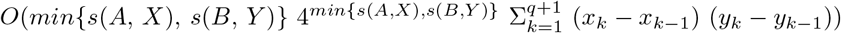 and 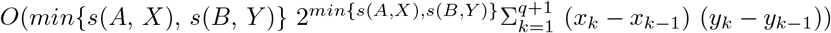, where *x*_0_ = *y*_0_ = 0, *x*_*q*+1_ = *m* + 1 and *y*_*q*+1_ = *n* + 1. As *X* as well as *Y* turns to have an increasing number of genes, both *min*{*s*(*A*, *X*), *s*(*B*, *Y*)} and 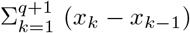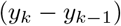 decrease. So there is an advantage of the algorithm: The algorithm can be dominated to take decreasing time and space to reach a solution by setting *X* as well as *Y* with increasing number of genes.

Using the algorithm for LCES-IG, we developed a software that have been ready for finding a longest common exemplar subsequence of two linear genomes *A* and *B*, which admits a subsequence identical to a given subsequence *X* of *A* as well as a given subsequence *Y* of *B*. Aiming at 23 human/gorilla chromosome pairs, we performed experiments where the algorithm was employed to quest for their longest common exemplar subsequences of annotated genes. By running the algorithm with *X* and *Y* as null, we obtained the longest common exemplar subsequences of 23 annotated gene summary pairs extracted from those 23 human/gorilla chromosome pairs.

In expectation of avail to find new conserved genes or new motifs in human as well as gorilla genomes, we performed experiments where the algorithm was employed to quest for longest common exemplar subsequences of *pseudo genes*, i.e., consecutive DNA subsequences that occurs in a human chromosome and its gorilla homologue, aiming at those 23 human/gorilla chromosome pairs. A modified version of SDquest [23] was used to identify the consecutive DNA subsequences in those 23 human/gorilla chromosome pairs as pseudo genes so that 23 pairs of pseudo gene sequences each of which will be called a *pseudo gene summary* were extracted from those 23 human/gorilla chromosome pairs.

For *X* and *Y* to be set with null, there are 20 pseudo gene summary pairs whose longest common exemplar subsequences have been found by the algorithm. The other 3 pseudo gene summary pairs were all verified solvable for the algorithm to reach their longest common exemplar subsequences that have to admit subsequences identical to given indexed gene subsequences *X* and *Y*. Positively, one is allowed to select more indexed genes in *X* as well as *Y* to make the algorithm take less time and space to reach a solution whose length can always come very close to a real longest common exemplar subsequence of the two given genomes. This implies it constantly true for the algorithm to reach an order conserved sequence of conserved consecutive DNA subsequences in a genome in as short a running time as required in practice.

In convenience for researchers to mine something from those longest common exemplar subsequences of pseudo genes, we performed experiments where we contrasted CES-7 and CES-16, the common exemplar subsequences of pseudo genes reached by the algorithm for the human chromosomes 7 and 16 and their gorilla homologues with the annotated genes in the human as well as gorilla chromosomes 7 and 16. One may be interested in the following statistics at least. There are 1 150 (resp. 2 353) pseudo genes in the human (resp. gorilla) chromosome map of CES-7 that do not overlap with any annotated gene. There are 1 552 (resp. 949) annotated genes in the human (resp. gorilla) chromosome 7 that overlap with some pseudo genes in the human chromosome map of CES-7, about 0.515 (resp. 0.659) times as many annotated genes as the human (resp. gorilla) chromosome 7 has. There are 528 (resp. 1 148) pseudo genes in the human (resp. gorilla) chromosome map of CES-16 that do not overlap with any annotated gene, meanwhile 1 420 (resp. 764) annotated genes in the human (resp. gorilla) chromosome 16 overlap with some pseudo genes, about 0.555 (resp. 0.668) times as many annotated genes as the human (resp. gorilla) chromosome 16 has.

## 2 Preliminaries

Let Σ be a set of *gene families*. An occurrence of a gene family in a genome is referred to as a *gene*. A sequence of genes is referred to as a *linear genome* on Σ, if all its genes are of the gene families in Σ. We usually mention a gene sequence instead of a linear genome to be on Σ if the gene sequence is unnecessary to represent a linear genome.

Let *π* = *π*[1] ... *π*[*m*] be a linear genome on Σ. We denote by *π*[*i*, *j*] the consecutive subsequence *π*[*i*] ... *π*[*j*] of *π* if 1 ≤ *i* ≤ *j* ≤ *m* or a null subsequence otherwise. A null (gene) sequence will be denoted as ””. Let *X* and *Y* be two gene sequences on Σ. We denote by *X || Y* the concatenation of *X* and *Y*. For example, *π*[1, 10] = *π*[1, 9] || *π*[10] = *π*[1] || *π*[2, 10] = *π*[1, 5] || *π*[6, 10].

There can be more than one gene of the same gene family in a linear genome or gene sequence. Two genes are referred to as *identical*, if they are of the same gene family. For a gene *π*[*i*] in *π*, we denote by *occ*(*π*, *π*[*i*]) the number of genes in *π* that are identical to *π*[*i*]. We refer to *j − i* for *i < j* as the *span* of *π*[*i*] and *π*[*j*] in *π*. The *span* of a linear genome refers to the maximum span of two identical genes in that linear genome. Let *π* = 1 4 3 2 4 1 5 2 6 for example. Then *occ*(*π*, 1) = *occ*(*π*, 2) = 2, *occ*(*π*, 3) = 1 and the span of *π* is 5, because *π*[1] and *π*[6] are identical and their span 5 is maximum over all two identical genes in *π*.

We denote it *x* = *y* for two genes *x* and *y* to be identical. Two gene sequences *X* = *X*[1, *p*] and *Y* = *Y* [1, *p*] are referred to as *identical*, if *X*[*i*] = *Y* [*i*] for *i* with 1 ≤ *i* ≤ *p*. A sequence of genes is referred to as a *subsequence* of a linear genome, if it can be obtained by deleting some genes (not necessarily consecutive) from the linear genome. A subsequence of a linear genome on Σ is referred to as an *exemplar subsequence* of the genome, if any two genes in this subsequence are not identical.

Let *π*[*x*_1_] ... *π*[*x*_*p*_] be an exemplar subsequence of *π* where 1 ≤ *x*_1_ *< ... < xp* ≤ *m*. Then *π*[*x*_*i*_] ≠ *π*[*x*_*j*_] for *x*_*i*_ ≠ *x*_*j*_, 1 ≤ *i*, *j* ≤ *p*. A *common exemplar subsequence* of two or more linear genomes refers to a gene sequence such that these linear genomes all admit an exemplar subsequence identical to the gene sequence. The *length* of a gene sequence refers to the number of genes in the gene sequence. For an arbitrary gene sequence *X*, we denote by |*X*| the length of *X*. A common exemplar subsequence of two or more linear genomes is referred to as *longest*, if it has no less genes than any common exemplar subsequence of these linear genomes.

## 3 LCES with indexed genes

A common exemplar gene subsequence of two or more linear genomes, if long enough, tends to reflect the structure similarity of these genomes so that, all genes in the common exemplar subsequence can be viewed as conserved. In practice, a certain number of conserved genes usually have been proven to occur in the given genomes in a consistent order. Then can we find a longest common exemplar subsequence of some linear genomes which admit proven-as-conserved genes in the same order as they are in the genomes? We describe this requirement in the form of a combinatorial optimization problem.

Let *G*_1_, *G*_2_, ..., *G*_*N*_ be linear genomes on Σ. Let *X*_*j*_ denote an exemplar subsequence of *G*_*j*_ for 1 ≤ *j* ≤ *N*. A common exemplar subsequence *C* of *G*_1_, *G*_2_, ..., *G*_*N*_ is referred to as *indexed by X*_*j*_, if there exists an exemplar subsequence of *G*_*j*_ identical to *C* such that *X*_*j*_ is a subsequence of it. The solution of the following problem is what we were just asked.

Instance: Linear genomes *G*_1_, *G*_2_, ..., *G*_*N*_; exemplar subsequences *X*_1_, *X*_2_, ..., *X*_*N*_ of *G*_1_, *G*_2_, ..., *G*_*N*_ respectively where *X*_1_ = *X*_2_ = ... = *X*_*N*_ and *N* ≥ 2.

Question: Find a longest common exemplar subsequence of *G*_1_, ..., *G*_*N*_ indexed by *X*_*j*_ for 1 ≤ *j ≤ N*.

We are going to mention this problem as LCES-IG for short and abbreviate by an LCES a longest common exemplar subsequence of some linear genomes.

In what follows of this section, we present a dynamic programming algorithm for LCES-IG whose instances are of two linear genomes and their respective exemplar subsequences. We represent an LCES-IG instance by two linear genomes *A* = *A*[1] ... *A*[*m*] and *B* = *B*[1] ... *B*[*n*] together with their respective exemplar subsequences *X* = *A*[*x*_1_] ... *A*[*x*_*q*_] and *Y* = *B*[*y*_1_] ... *B*[*y*_*q*_] where *X* = *Y*, 1 ≤ *x*_*i*_ < *x*_*j*_ ≤ *m*, 1 ≤ *y*_*i*_ < *y*_*j*_ ≤ *n* for 1 ≤ *i < j ≤ q*. A gene in *A* (resp. *B*), if identical to a gene in *X* (resp. *Y*) and other than the gene, cannot occur in any common exemplar subsequence indexed by *X* (resp. *Y*). Thus *A* (resp. *B*) is always assumed with no other gene identical to *A*[*x*_*k*_] (resp. *B*[*y*_*k*_]) than *A*[*x*_*k*_] (resp. *B*[*y*_*k*_]) for 1 ≤ *k* ≤ *q*.

### 3.1 Common exemplar subsequences

For *i* and *j* with 0 ≤ *i ≤ m* and 0 ≤ *j ≤ n*, let *C*(*i*, *j*) denote the set of all common exemplar subsequences of *A*[1, *i*] and *B*[1, *j*]. A member *C ∈ C*(*i*, *j*) is referred to as an *extension* of a member *C′ ∈ C*(*i − x*, *j − y*) for *x ≥* 1 and *y ≥* 1 if there exists a common exemplar subsequence *C′′* of *A*[*i − x* + 1,*i*] and *B*[*j − y* + 1, *j*] such that *C* = *C′ || C′′*. Only when *i ≥ x*_*k*_ and *j ≥ y*_*k*_, can someone in *C*(*i*, *j*) be indexed by *X*[1, *k*] as well as *Y* [1, *k*]. The following lemma concentrates more attention on the value intervals of *i* and *j* in which a dynamic programming should maintain in *C*(*i, j*) those members indexed by *X*[1*, k*] as well as *Y* [1*, k*].

#### Lemma 1.

*If for i ≥ x*_*k*_ (*resp. j ≥ y*_*k*_) *where* 1 ≤ *k ≤ q, a member in C*(*i, j*) *fails to be indexed by X*[1*, k*] *as well as Y* [1*, k*]*, then any extension of the member cannot be indexed by X as well as Y.*

*Proof.* Let *C*_1_ ∈ *C*(*i, j*), *C* ∈ *C*(*m, n*). If *C* is an extension of *C*_1_, then there is a common exemplar subsequence *C*_2_ of *A*[*i* + 1, *m*] and *B*[*j* + 1, *n*], such that *C* = *C*_1_ ∥ *C*_2_.

It follows from *occ*(*A*, *A*[*x*_*p*_]) = 1 for 1 ≤ *p* ≤ *k* that *C*_2_ cannot be indexed by *X*[*p*] for 1 ≤ *p* ≤ *k*. If *C*_1_ fails to be indexed by *X*[1, *k*] as well as *Y* [1, *k*], so does *C*.

It follows by Lemma 1 that if *i ≥ x*_*k*_ and *j ≥ y*_*k*_, anyone in *C*(*i, j*) can be given up to maintain for extension if it fails to be indexed by *X*[1, *k*] as well as *Y* [1, *k*]. This implies that if *i ≥ x*_*k*+1_ and *j ≥ y*_*k*+1_ for 1 ≥ *k* + 1 ≥ *q*, then there is no need to maintain for extension any one in *C*(*i*, *j*) that is indexed by *X*[1, *k*] as well as *Y*[1, *k*] instead of *X*[1, *k* + 1] as well as *Y*[1, *k* + 1]. On the other hand, if *i* ≥ *x*_*k*+1_ and *j < y*_*k*+1_ (resp. *i < x*_*k*+1_ and *j ≥ y*_*k*+1_) for 1 ≤ *k* + 1 ≤ *q*, then it follows by Lemma 1 that anyone in *C*(*i*, *j*) can be given up to maintain for extension. Thus only when *i* and *j* fall in such intervals as *x*_*k*_ ≤ *i < x*_*k*+1_ and *y*_*k*_ ≤ *j < y*_*k*+1_, is it necessary to maintain for extension those common exemplar subsequences in *C*(*i*, *j*) indexed by *X*[1, *k*] as well as *Y* [1, *k*],

Let *x*_0_ = *y*_0_ = 0, *x*_*q*+1_ = *m* + 1, *y*_*q*+1_ = *n* + 1. We treat *A*[*x*_0_] and *A*[*x*_*q*+1_] (resp. *B*[*y*_0_] and *B*[*y*_*q*+1_]) to be null genes and reexpress *A* and *B* by *A* = *A*[0] *A*[1] ... *A*[*m*] *A*[*m* + 1] and *B* = *B*[0] *B*[1] ... *B*[*n*] *B*[*n* + 1]. Let for *i* and *j* with *x*_*k*_ ≤ *i < x*_*k*+1_ and *y*_*k*_ ≤ *j < y*_*k*+1_ where 0 ≤ *k ≤ q*, 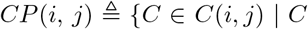 is indexed by *X*[1*, k*] as well as *Y* [1*, k*]. Then in *CP*(*m*, *n*), there must exist an LCES of *A* and *B* indexed by *X* as well as *Y*.

For *i* and *j* with *x*_*k*_ ≤ *i < x*_*k*+1_ and *y*_*k*_ ≤ *j < y*_*k*+1_ (0 ≤ *k* ≤ *q*), we focus on how get *CP*(*i, j*). Since *A*[1, 0] = ”” and *B*[1, 0] = ””, we set *CP* (*x*_0_, *y*_0_) = *CP*(0, 0) = {””}. The following lemma is in terms of how to get *CP* (*x*_*k*_, *y*_*k*_) for *k* ≥ 1.

#### Lemma 2.

*For k with* 1 ≤ *k* ≤ *q, a common exemplar subsequence C* ∈ *C*(*x*_*k*_, *y*_*k*_) *is indexed by X*[1*, k*] *as well as Y* [1*, k*]*, if and only if there exists C′* ∈ *C*(*x*_*k*_ − 1*, y*_*k*_ − 1]) *indexed by X*[1*, k* − 1] *as well as Y* [1*, k* − 1] *such that C* = *C′* ∥ *A*[*x*_*k*_].

*Proof.* **If**: Let *C′ ∈ C*(*x*_*k*_ − 1, *y*_*k*_ − 1). It follows from *occ*(*A*, *A*[*x*_*k*_]) = *occ*(*B*, *B*[*y*_*k*_]) = 1 that no gene identical to *A*[*x*_*k*_] or *B*[*y*_*k*_] can occur in *C′*. It follows from *A*[*x*_*k*_] = *B*[*y*_*k*_] that *C′* ∥ *A*[*x*_*k*_] ∈ *C*(*x*_*k*_, *y*_*k*_) and if *C′* is indexed by *X*[1, *k* − 1] as well as *Y*[1, *k* − 1], then *C′* ∥ *A*[*x*_*k*_] is indexed by *X*[1, *k*] as well as *Y* [1, *k*].

**Only if**: Let *C* ∈ *C*(*x*_*k*_, *y*_*k*_) be indexed by *X*[1, *k*] as well as *Y* [1, *k*]. Since *A*[*x*_*k*_] = *B*[*y*_*k*_] and *occ*(*A*, *A*[*x*_*k*_]) = *occ*(*B*, *B*[*y*_*k*_]) = 1, *C* can be expressed as *C′* ∥ *A*[*x*_*k*_] where *C′* ∈ *C*(*x*_*k*_ − 1, *y*_*k*_ − 1). Since *C* is indexed by *X*[1, *k*] as well as *Y* [1, *k*], *C′* must be indexed by *X*[1, *k* − 1] as well as *Y* [1, *k* − 1].

Let 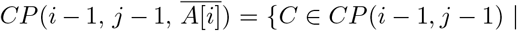 no identical gene to *A*[*i*] occurs in *C*} and 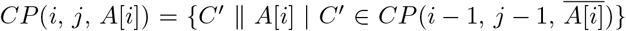. Since *occ*(*A*, *A*[*x*_*k*_]) = 1, 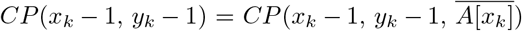 for 1 ≤ *k* ≤ *q*. If *k* ≥ 1, it follows by Lemma 2 that *CP*(*x*_*k*_, *y*_*k*_) = {*C* ∥ *A*[*x*_*k*_] | *C* ∈ *CP*(*x*_*k*_ − 1, *y*_*k*_ − 1)} = *CP*(*x*_*k*_, *y*_*k*_, *A*[*x*_*k*_]). It remains to argue the way of how to get *CP*(*i*, *j*) from *CP*(*i*, *j* − 1), *CP*(*i* − 1, *j*) and *CP*(*i* − 1, *j* − 1) for *i* and *j* with *x*_*k*_ ≤ *i < x*_*k*+1_ and *y*_*k*_ ≤ *j < y*_*k*+1_ (0 ≤ *k* ≤ *q*) other than *i* = *x*_*k*_ and *j* = *y*_*k*_.

(1)If *x*_*k*_ < *i < x*_*k*+1_ and *y*_*k*_ < *j < y*_*k*+1_, then each member in *CP*(*i* − 1, *j*) ∪ *CP*(*i*, *j* − 1) ∪ *CP*(*i* − 1, *j* − 1) is indexed by *X*[1, *k*] as well as *Y*[1, *k*], thus *CP*(*i* − 1, *j*) ∪ *CP*(*i*, *j* − 1) ∪ *CP*(*i* − 1, *j* − 1) ⊆ *CP*(*i*, *j*). Let *C* = *C*[1] *C*[2] ... *C*[*p*] = *A*[*a*_1_] *A*[*a*_2_] ... *A*[*a*_*p*_] = *B*[*b*_1_] *B*[*b*_2_] ... *B*[*b*_*p*_] denote an arbitrary common exemplar subsequence of *A*[1, *i*] and *B*[1, *j*] indexed by *X*[1, *k*] as well as *Y*[1, *k*].

(1.1)If *A*[*i*] ≠ *B*[*j*], then *a*_*p*_ ≠ *i* or *b*_*p*_ ≠ *j*. This implies *C* ∈ *CP*(*i* − 1, *j*) or *C* ∈ *CP*(*i*, *j* − 1). Thus *CP*(*i*, *j*) ⊆ *CP*(*i* − 1, *j*) ∪ *CP*(*i*, *j* − 1). It follows from *CP*(*i* − 1, *j*) ∪ *CP*(*i*, *j* − 1) ⊆ *CP*(*i*, *j*) that *CP*(*i*, *j*) = *CP*(*i* − 1, *j*) ∪ *CP*(*i*, *j* − 1).

(1.2)Assume *A*[*i*] = *B*[*j*]. It follows from *CP*(*i* − 1, *j* − 1) ⊆ *CP*(*i*, *j*) that *CP*(*i*, *j*, *A*[*i*]) ⊆ *CP*(*i*, *j*). If a_p_ ≠ *i* or b_p_ ≠ *j*, then *C* ∈ *CP*(*i* − 1, *j*) or *C* ∈ *CP*(*i*, *j* − 1). Otherwise, it follows from *a*_*p*_ = *i* and *b*_*p*_ = *j* that *C*[*p*] = *A*[*i*] and 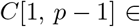 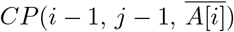. Thus *C* ∈ *CP*(*i*, *j*, *A*[*i*]). To sum up, *CP*(*i*, *j*) ⊆ *CP*(*i* − 1, *j*) ∪ *CP*(*i*, *j* − 1) ∪ *CP*(*i*, *j*, *A*[*i*]). It follows from *CP*(*i* − 1, *j*) ∪ *CP*(*i*, *j* − 1) ∪ *CP*(*i*, *j*, *A*[*i*]) ⊆ *CP*(*i*, *j*) that *CP*(*i*, *j*) = *CP*(*i* − 1, *j*) ∪ *CP*(*i*, *j* − 1) ∪ *CP*(*i*, *j*, *A*[*i*]).

(2)If *i* = *x*_*k*_ and *y*_*k*_ < *j < y*_*k*+1_, then it follows from *occ*(*B, B*[*y*_*k*_]) = 1 and *A*[*x*_*k*_] = *B*[*y*_*k*_] that *A*[*i*] ≠ *B*[*j*]. In this case, we have *CP*(*i*, *j*) ⊆ *CP*(*i* − 1, *j*) ∪ *CP*(*i*, *j* − 1), that can be argued true in the same way as in case (1.1). Since *i* = *x*_*k*_, no member in *CP*(*i* − 1, *j*) can be indexed by *X*[1, *k*] as well as *Y* [1, *k*], which means *CP*(*i, j*) ⊆ *CP*(*i*, *j* − 1). Since each member in *CP*(*i*, *j* − 1) is indexed by *X*[1, *k*] as well as *Y* [1, *k*], *CP*(*i*, *j*) = *CP*(*i*, *j* − 1).

(3)If *j* = *y*_*k*_ and *x*_*k*_ < *i < x*_*k*+1_, then *CP*(*i*, *j*) = *CP*(*i* − 1, *j*) for the same reason. To sum up, *CP*(*i*, *j*) can be computed recursively by Formula (1) for *i* and *j* with *i* + *j >* 0, *x*_*k*_ ≤ *i < x*_*k*+1_ and *y*_*k*_ ≤ *j < y*_*k*+1_ (0 ≤ *k* ≤ *q*).

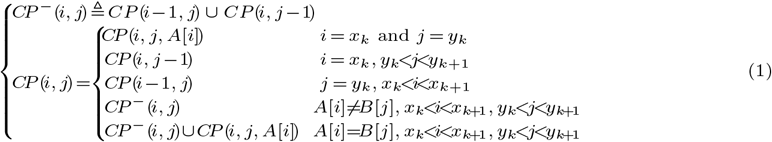

### 3.2 Confused gene family set

Let *C* be a common exemplar subsequence of *A*[1, *i*] and *B*[1, *j*]. A gene family that occurs in *C* is referred to as *confused*, if it also occurs in *A*[*i* + 1, *m*] as well as *B*[*j* + 1, *n*]. The *confused gene family set* of *C* refers to the set of all confused gene families that occur in *C*, and will be denoted as *f*(*i, j*, *C*). If two common exemplar subsequences in *CP*(*i*, *j*) admit the same confused gene family set, then one of them can be given up for extension.

#### Lemma 3.

*Let for i and j with x*_*k*_ ≤ *i < x*_*k*+1_ *and y*_*k*_ ≤ *j < y*_*k*+1_ *where* 0 ≤ *k* ≤ *q, C*_1_ ∈ *CP*(*i, j*) *and C*_2_ ∈ *CP*(*i, j*)*. If f*(*i, j, C*_1_) = *f*(*i, j, C*_2_) *and* |*C*_1_| ≥ |*C*_2_|*, then in CP*(*m, n*)*, a longest extension of C*_1_ *is no shorter than any extension of C*_2_.

*Proof.* Let *C* be a longest extension of *C*_2_ in *CP*(*m*, *n*). Then there exists a common exemplar subsequence of *A*[*i* + 1, *m*] and *B*[*j* + 1, *n*] indexed by *X*[*k* + 1, *q*] as well as *Y*[*k* + 1, *q*], say *C*_3_ such that *C* = *C*_2_ ∥ *C*_3_. Then it follows from *f*(*i*, *j*, *C*_1_) = *f*(*i*, *j*, *C*_2_) and *C*_1_ is indexed by *X*[1, *k*] as well as *Y*[1, *k*] that *C*_1_ ∥ *C*_3_ is a common exemplar subsequence of *A* and *B* indexed by *X* as well as *Y* that is an extension of *C*_1_ in *C*(*m, n*). Since |*C*_1_| ≥ |*C*_2_|, |*C*_1_ ∥ *C*_3_| ≥ |*C*|. The lemma follows from that the longest extension of *C*_1_ in *C*(*m, n*) has no less genes than |*C*_1_ ∥ *C*_3_|.

By Lemma 3, to get an LCES of *A* and *B* indexed by *X* as well as *Y*, it suffices for the dynamic programming to maintain a subset of *CP*(*i*, *j*) whose common exemplar subsequences admit mutually distinct confused gene family sets. A subset of *CP*(*i*, *j*) is referred to as *representative*, if there exists a mapping from *CP*(*i, j*) to the subset, such that each member in *CP*(*i, j*) admits the same confused gene family set as its image in the subset, and have no more genes than its image in the subset. A representative subset of *CP*(*i*, *j*) is referred to as *minimum*, if any two members in it do not admit the same confused gene family set. Let *CFP*(*i*, *j*) denote an arbitrary minimum representative subset of *CP*(*i, j*). An LCES of *A* and *B* indexed by *X* as well as *Y* must occur in a minimum representative subset of *CP*(*m*, *n*). Instead of getting a minimum representative subset of *CP*(*i*, *j*) from *CP*(*i, j*), we pursue to arrive at a minimum representative subset of *CP*(*i*, *j*) from *CFP*(*i* − 1, *j*), *CFP*(*i*, *j* − 1) and *CFP*(*i* − 1, *j* − 1).

It follows from *CP*(0, 0) = {””} that *CFP*(0, 0) = {””}. Then for *i* and *j* with *i* + *j >* 0, *x*_*k*_ ≤ *i < xk*+1 and *y*_*k*_ ≤ *j < y*_*k*+1_ (0 ≤ *k* ≤ *q*), we face with the task of getting a minimum representative subset of *CP*(*i, j*) from *CFP*(*i* − 1, *j*), *CFP*(*i*, *j* − 1) and *CFP*(*i* − 1, *j* − 1).

Let 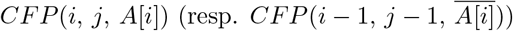 denote a minimum representative subset of 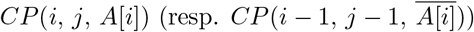. If *A*[*i*] = *B*[*j*] for *i* and *j* with *i* + *j >* 0, *x*_*k*_ ≤ *i* < *x*_*k*+1_ and *y*_*k*_ ≤ *j* < *y*_*k*+1_ (0 ≤ *k ≤ q*), a minimum representative subset of *CP*(*i*, *j*, *A*[*i*]) can be identified by the following lemma.

#### Lemma 4.

*If A*[*i*] = *B*[*j*] *for i and j with i* + *j >* 0*, x*_*k*_ ≤ *i* < *x*_*k*+1_ *and y*_*k*_ ≤ *j* < *y*_*k*+1_ (0 ≤ *k ≤ q*)*, then 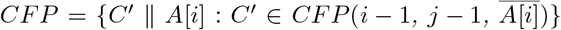 is a minimum representative subset of CP*(*i, j, A*[*i*]).

*Proof.* A common exemplar subsequence in *CP*(*i*, *j*, *A*[*i*]) always takes the form of *C′ || A*[*i*] where 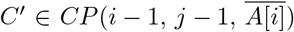. It follows from 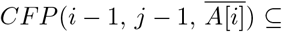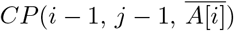 that *CFP ⊆ CP*(*i*, *j*, *A*[*i*]). Then we argue for *CFP* to be representative in the following two aspects.

(1)Let 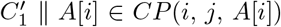 where 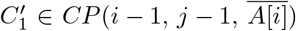. Then since 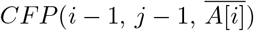 is representative, there exists a 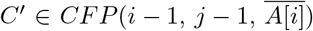 with 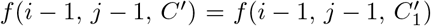. Then 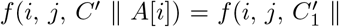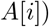.

(2)Let *C′* ∥ *A*[*i*] ∈ *CFP*, 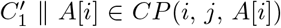. If 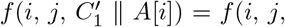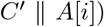, then since 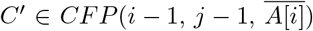,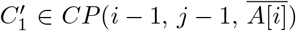, 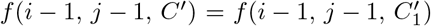. It follows from 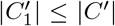 that 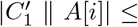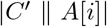.

The reason why *CFP* is minimum over all representative subsets of *CP*(*i*, *j*, *A*[*i*]) lies in that 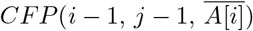 is minimum over all those representative subsets of 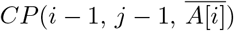.

For *i* and *j* with *i* + *j >* 0, *x*_*k*_ ≤ *i < x*_*k*+1_ and *y*_*k*_ ≤ *j < y*_*k*+1_ (0 ≤ *k ≤ q*), it follows from Formula (1) that a minimum representative subset of *CP*(*i*, *j*) is included in a subset of *CFP*(*i* − 1, *j*) ∪ *CFP*(*i*, *j* − 1) ∪ *CFP*(*i*, *j*, *A*[*i*]) as described in the following formula where 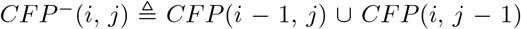.

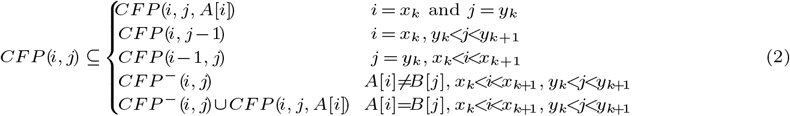

Let *CP*(*i*, *j*, *CFP*(*i*, *j −* 1)) = *C ∈ CP*(*i*, *j*) *C ∈ CFP*(*i*, *j −* 1), *CP*(*i*, *j*, *CFP*(*i* − 1, *j*)) = *C ∈ CP*(*i*, *j*) *C ∈ CFP*(*i −* 1, *j*). A minimum representative subset of *CP*(*i, j*) can be extracted by examining every two members in a subset of *CP*(*i*, *j*, *CFP*(*i*, *j −* 1)) ∪ *CP*(*i*, *j*, *CFP*(*i* − 1, *j*)) ∪ *CFP*(*i*, *j*, *A*[*i*]) for if they admit the same confused gene family set and if yes, throwing away that one no longer than the other. Thus, to get a minimum representative subset of *CP*(*i, j*), it remains to calculate the confused gene family set of *C* = *C′* ∥ *A*[*i*] ∈ *CFP*(*i*, *j*, *A*[*i*]) where 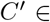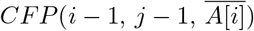 and the confused gene family set of *C* ∈ *CP*(*i*, *j*, *CFP*(*i*, *j* − 1)) ∪ *CP*(*i*, *j*, *CFP*(*i* − 1, *j*)) for *i* and *j* with *i* + *j >* 0, *x*_*k*_ ≤ *i < x*_*k*+1_ and *y*_*k*_ ≤ *j < y*_*k*+1_ (0 ≤ *k ≤ q*).

(1)If *C* = *C′ ∥ A*[*i*] ∈ *CFP*(*i*, *j*, *A*[*i*]) where 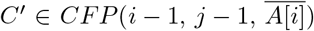, a gene family in *f*(*i −* 1, *j −* 1, *C′*) must be confused where it is treated as what occurs in *C′ ∥ A*[*i*] *CP*(*i*, *j*). On the other hand, no more gene family than that of *A*[*i*] can become confused where it is treated as what occurs in *C*. To get *f*(*i*, *j*, *C*), it suffices to do according to the following two sub cases.

(1.1) If no identical gene to *A*[*i*] occurs in *A*[*i* + 1, *m*] or *B*[*j* + 1, *n*], then the gene family of *A*[*i*] will go outside *f*(*i*, *j*, *C′ A*[*i*]).

(1.2) If an identical gene to *A*[*i*] occurs in both *A*[*i* + 1, *m*] and *B*[*j* + 1, *n*], then the gene family of *A*[*i*] must fall in *f*(*i*, *j*, *C′ A*[*i*]).

Formally, the confused gene family set of *C* can be got as follows.

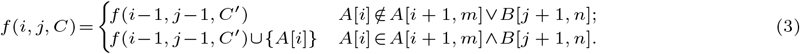

(2) Assume *C ∈ CP*(*i*, *j*, *CFP*(*i*, *j −* 1)) ∪ *CP*(*i*, *j*, *CFP*(*i −* 1, *j*)). If a confused gene family in *f*(*i −* 1, *j*, *C*) (resp. *f*(*i*, *j* − 1, *C*)) is other than that of *A*[*i*] (resp. *B*[*j*]), it keeps confused where it is treated as what occurs in *C ∈ CP*(*i*, *j*). No more gene family than those in *f*(*i −* 1, *j*, *C*) as well as *f*(*i*, *j −* 1, *C*) can become confused where it is treated as what occurs in *C ∈ CP*(*i*, *j*). Thus to get the confused gene family set of *C ∈ CP*(*i, j*), it suffices to decide whether the gene family of *A*[*i*] (resp. *B*[*j*]) in *f*(*i* − 1, *j*, *C*) (resp. *f*(*i*, *j* − 1, *C*)), is confused or not where it is treated as what occurs in *C ∈ CP*(*i*, *j*). This can be argued in two subcases.

(2.1) If no identical gene to *A*[*i*] (resp. *B*[*j*]) occurs in *A*[*i* + 1, *m*] or *B*[*j* + 1, *n*], then the gene family of *A*[*i*] (resp. *B*[*j*]) goes out of confused where we treat it as what occurs in *C ∈ CP*(*i*, *j*).

(2.2) Otherwise, the gene family of *A*[*i*] (resp. *B*[*j*]) remains confused where we treat it as what occurs in *C ∈ CP*(*i*, *j*).

In summary of (2.1) and (2.2), *f*(*i*, *j*, *C*) can be got by the following formula.

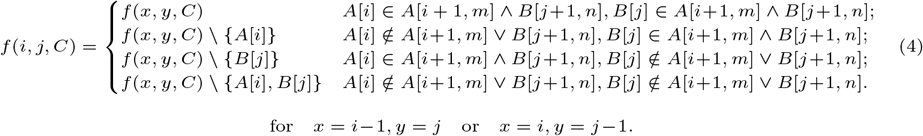

### 3.3 The algorithm

Aiming for the dynamic programming to use less storage space, we choose to maintain the confused gene family sets and lengths of those members in *CFP*(*i*, *j*) instead of *CFP*(*i, j*) itself. For *CX ⊆ CP*(*i*, *j*), we refer to the multi set with those confused gene family sets of all members in *CX* as the *confused gene family set collection* of *CX*.

Let *FP*(*i*, *j*) denote the confused gene family set collection of *CFP*(*i*, *j*). Since *CFP*(*i*, *j*) is minimum, *f*(*i*, *j*, *C*) for *C* ∈ *CFP*(*i, j*) is a 1-1 mapping from *CFP*(*i*, *j*) to *FP*(*i*, *j*). Since *CFP*(0, 0) = {””}, we set *FP*(0, 0) = {∅}. For *i* and *j* with *i* + *j >* 0, *x*_*k*_ ≤ *i* < *x*_*k*+1_ and *y*_*k*_ ≤ *j* < *y*_*k*+1_ (0 ≤ *k ≤ q*), we argue that *FP*(*i*, *j*) can be arrived at from *FP*(*i* − 1, *j*), *FP*(*i*, *j* − 1) and *FP*(*i* − 1, *j* − 1).

Let respectively, *FP*(*i*, *j*, *CFP*(*i* − 1, *j*)) and *FP*(*i*, *j*, *CFP*(*i*, *j* − 1)) denote the confused gene family set collections of *CP*(*i*, *j*, *CFP*(*i* − 1, *j*)) and *CP*(*i*, *j*, *CFP*(*i*, *j* − 1)). Then all members in *FP*(*i*, *j*, *CFP*(*i* − 1, *j*)) ∪ *FP*(*i*, *j*, *CF*(*i*, *j* − 1)) can be computed from those in *FP*(*i* − 1, *j*) ∪ *FP*(*i*, *j* − 1) by Formula (4). Let *FP*(*i*, *j*, *A*[*i*]) denote the confused gene family set collection of *CFP*(*i*, *j*, *A*[*i*]). Then all members in *FP*(*i*, *j*, *A*[*i*]) can be computed from those in *FP*(*i* 1, *j* 1) by Formula (3).

To extract *FP*(*i*, *j*), it needs help of the lengths of those members in *CP*(*i*, *j*, *CFP*(*i* − 1, *j*)), *CP*(*i*, *j*, *CFP*(*i*, *j* − 1)) and *CFP*(*i*, *j*, *A*[*i*]). For *f* = *f*(*i*, *j*, *C*) where *C* ∈ *CP*(*i*, *j*), we refer to the length of *C* as the *CES length* of *f* and denote by *L*(*f*) the CES length of *f*. Since *CFP*(0, 0) = {””}, the CES length of the unique member in *FP*(0, 0) should be assigned with 0. For *i* and *j* with *i* + *j >* 0, *x*_*k*_ ≤ *i < x*_*k*+1_ and *y*_*k*_ ≤ *j < y*_*k*+1_ (0 ≤ *k ≤ q*), the CES length of an arbitrary confused gene family set *f ∈ FP*(*i*, *j*, *CFP*(*i* − 1*, j*)) ∪ *FP*(*i*, *j*, *CFP*(*i, j* − 1)) ∪ *FP*(*i*, *j*, *A*[*i*]) can be computed recursively by Formula (5).

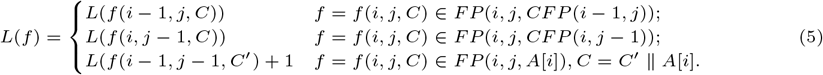

For *CX ⊆ CP*(*i*, *j*), let *F*[*CX*] denote the confused gene family set collection of *CX*. We set a subroutine named as *U*(*F*[*CX*]) to extract the confused gene family set collection of a minimum representative subset of *CX*. To get *U*(*F*[*CX*]) from *F*[*CX*], it suffices to examine every two members in *F*[*CX*] for if they are equal to each other, and if yes, removing from them that one whose CES length is no larger than the other. For *i* and *j* with *i* + *j >* 0, *x*_*k*_ ≤ *i < x*_*k*+1_ and *y*_*k*_ ≤ *j < y*_*k*+1_ (0 ≤ *k ≤ q*), since all member in *FP*(*i*, *j*, *CFP*(*i −* 1*, j*)) ∪ *FP*(*i*, *j*, *CFP*(*i, j* − 1)) ∪ *FP*(*i*, *j*, *A*[*i*]) is accompanied with a CES length, *FP*(*i*, *j*) can be arrived at recursively by Formula (6).

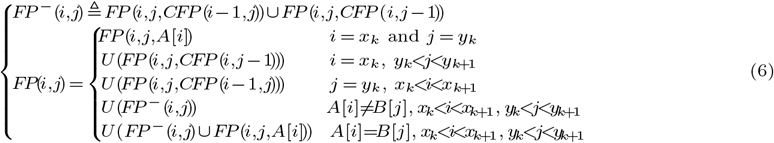

This leads to a dynamic programming based algorithm to get *FP*(*m, n*). Since all members in *CP*(*m, n*) admit empty confused gene family sets, there exists a unique empty confused gene family set in *FP*(*m, n*) that is of an LCES of *A* and *B* indexed by *X* as well as *Y*. By tracing back from *FP*(*m*,*n*) to *FP*(*i*, *j*) where *i* = 0 or *j* = 0, an LCES of *A* and *B* indexed by *X* as well as *Y* can be got. In Algorithm 1, we present pseudo codes of how to get an LCES of *A* and *B* indexed by *X* as well as *Y*. If we set *X* as well as *Y* as the null gene sequence ””, the solution of Index-LCES(*A*, *B*, *X*, *Y*) is an LCES of *A* and *B*.

**Algorithm 1.**
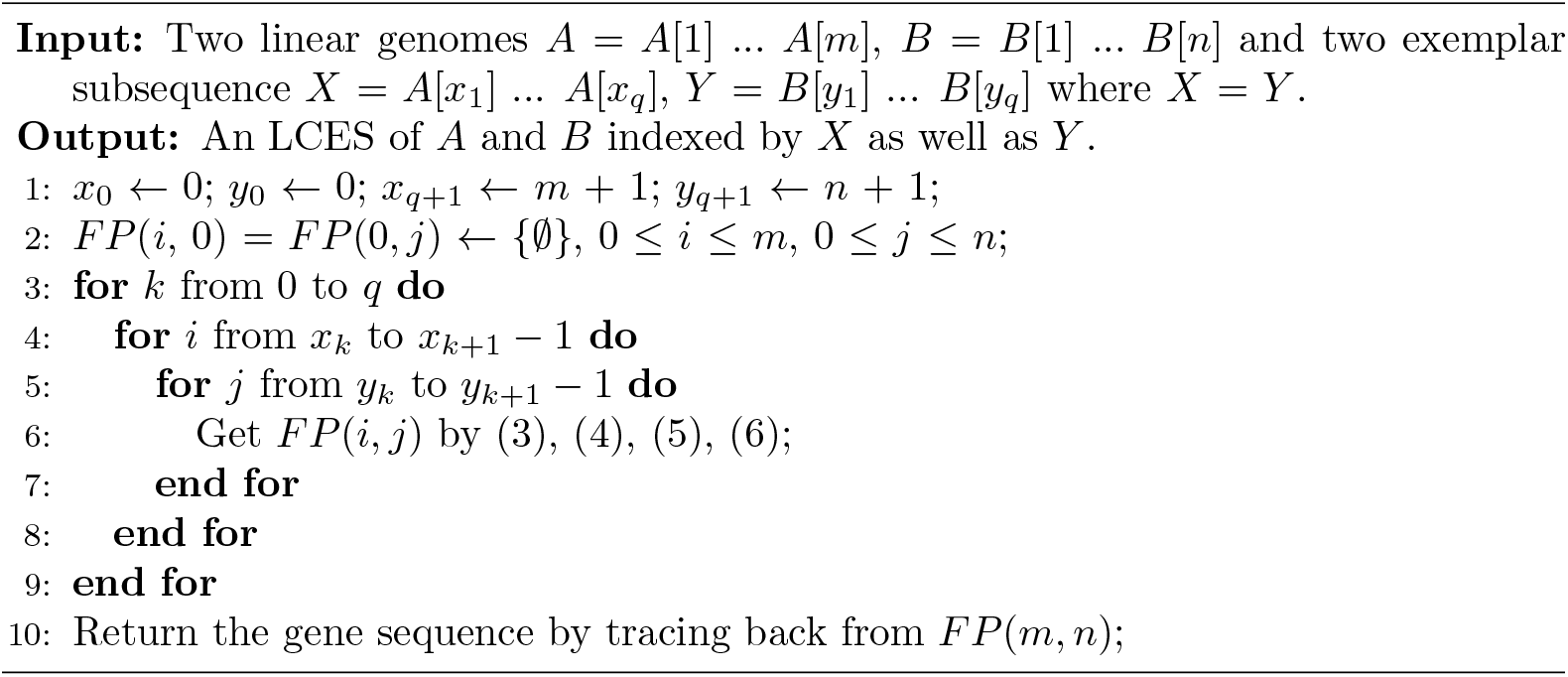
Index-LCES(*A, B, X, Y*)

### 3.4 The algorithm complexity

The span of two identical genes in a linear genome has usually been used as a parameter to design efficient algorithms [11] [14]. There have been approaches on comparing human and mouse genomes which provide hints for identical genes in a genome to occur in limited spans [24]. Since so, let *s*(*A*, *X*) (resp. *s*(*B*,*Y*)) denote the span of *A*(resp. *B*) in exclusion of those genes each of which is identical to someone in *X*(resp. *Y*), 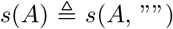 and 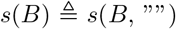. It follow from *occ*(*A*, *A*[*x*_*k*_]) = *occ*(*B*, *B*[*y*_*k*_]) = 1 for *k* with 1 ≤ *k ≤ q* that *s*(*A*) = *s*(*A*, *X*) and *s*(*B*) = *s*(*B*, *Y*). A confused gene family set collection that happens in LCES(*A*, *B*, *X*, *Y*) is bounded in size as follows.

#### Lemma 5.

*For i and j with x*_*k*_ ≤ *i < x*_*k*+1_ *and y*_*k*_ ≤ *j < y*_*k*+1_ (0 ≤ *k* ≤ *q*), |*FP*(*i, j*)| ≤ 2^*min{s*(*A*)*,s*(*B*)}^.

*Proof.* Without loss of generality, let *s*(*A*) = *min*{*s*(*A*)*, s*(*B*)}. Then there are at most *s*(*A*) gene families in both *A*[1, *i*] and *A*[*i* + 1, *m*] for any *i* with 0 ≤ *i ≤ m*. A confused gene family of an arbitrary member in *CP*(*i, j*) must occur in both *A*[*i* + 1,*m*] and *B*[*j* + 1, *n*]. Since at most *s*(*A*) gene families can occur in both *A*[1, *i*] and *A*[*i* + 1, *m*], an arbitrary member in *CP*(*i*, *j*) can admit a confused gene family set of at most *s*(*A*) gene families. The lemma follows from that those confused gene family sets in *FP*(*i*, *j*) are mutually different.

Let *x*_0_ = *y*_0_ = 0, *x*_*q*+1_ = *m* + 1 and *y*_*q*+1_ = *n* + 1. It is necessary for LCES(*A*, *B*, *X*, *Y*) to compute and maintain 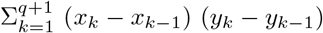 confused gene family set collections. By Lemma 5, every member in *FP*(*i*, *j*) has at most 2^*min{s*(*A*)*,s*(*B*)}^ gene families. Thus the space complexity of LCES(*A*, *B*, *X*, *Y*) is 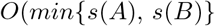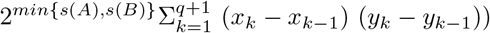. Subsequently, the time complexity of the algorithm is 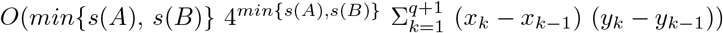. If one would like to assume *occ*(*A*, *A*[*x*_*k*_]) = *occ*(*B*, *B*[*y*_*k*_]) ≠ 1 for some *k*, then the time complexity and space complexity of the algorithm remain as above with no other exception than the substitution of *s*(*A*) and *s*(*B*) with *s*(*A*, *X*) and *s*(*B*, *Y*).

#### Theorem 6.

*The algorithm LCES*(*A, B, X, Y*) *can get an LCES of A and B indexed by X as well Y in* 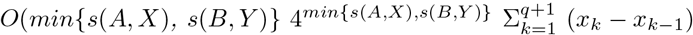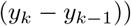 *time and* 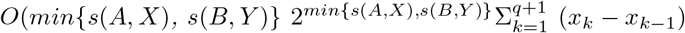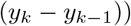 *space.*

## 4 LCES of more than two linear genomes

Let for *N ≥* 2, *G*_1_, *G*_2_, ..., *G*_*N*_ be *N* linear genomes on Σ whose lengths are *n*_1_, *n*_2_, ..., *n*_*N*_ respectively. Let 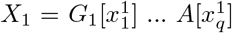, 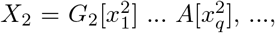, ..., and 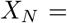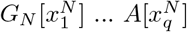 denote the respective exemplar subsequences of *G*_1_, *G*_2_, ..., *G*_*N*_ where *X*_1_ = *X*_2_ = ... = *X*_*N*_. Let *s* = *s*(*G*_1_) = *min*{*s*(*G*_1_), ..., (*G*_*N*_)}. Let *CP*(*i*_1_, *i*_2_, ..., *i*_*N*_) denote the set of all common exemplar subsequences of *G*_1_[1, *i*_1_], *G*_2_[1, *i*_2_], ..., *G*_*N*_ [1, *i*_*N*_] indexed by *X*_*j*_ for 1 ≤ *j* ≤ *N*.

For *C* ∈ *CP*(*i*_1_, *i*_2_, ..., *i*_*N*_), a gene family that occurs in *C* is referred to as *confused*, if it also occurs in *G*_1_[*i*_1_ + 1, *n*_1_], *G*_2_[*i*_2_ + 1, *n*_2_], ..., *G*_*N*_ [*i*_*N*_ + 1, *n*_*N*_]. The *confused gene family set* of *C* refers to the set of all confused gene families that occur in *C*. Let *CFP*(*i*_1_, *i*_2_, ..., *i*_*N*_) be an arbitrary minimum representative subset of *CP*(*i*_1_, *i*_2_, ..., *i*_*N*_) and *FP*(*i*_1_, *i*_2_, ..., *i*_*N*_) a confused gene family set collection of *CFP*(*i*_1_, *i*_2_, ..., *i*_*N*_).

Since *G*_*j*_[1,0] = ”” for 1 ≤ *j* ≤ *N*, *FP*(0, 0, ..., 0) = {∅}. Then, let 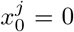 and 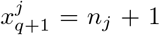 with 1 ≤ *j* ≤ *N*, for *i*_1_ + *i*_2_ + ... + *i*_*N*_ > 0 and 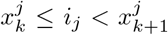 where 1 ≤ *j* ≤ *N* and 0 ≤ *k* ≤ *q*, the confused gene family set collection *FP*(*i*_1_, *i*_2_, ..., *i*_*N*_) can be arrived at from all confused gene family set collections in the form of *FP*(*x*_1_, *x*_2_, ..., *x*_*k*_) for *x*_*j*_ ∈ {*i*_*j*_ − 1, *i*_*j*_} (1 ≤ *j* ≤ *N*) except *FP*(*i*_1_, *i*_2_, ..., *i*_*N*_). To get *FP*(*i*_1_, *i*_2_, ..., *i*_*N*_), it has to access 2^*N*^ − 1 confused gene family set collections. There are at most *s* gene families that can occur in both *G*_1_[1, *i*] and *G*_1_[*i* + 1, *n*_1_] for any *i* with 1 ≤ *i ≤ n*_1_. Then a member in *CP*(*i*_1_, *i*_2_, ..., *i*_*N*_) admits a confused gene family set of at most *s* gene families and there are at most 2^*s*^ members in *FP*(*i*_1_, *i*_2_, ..., *i*_*N*_). In order to go from *FP*(0, 0, ..., 0) to *FP*(*n*_1_, *n*_2_, ..., *n*_*k*_), there are 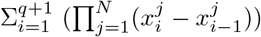 confused gene family set collections to be involved into computation, where 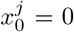 and 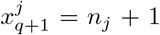 for 1 ≤ *j* ≤ *N*.

So, it can take 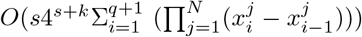 time and 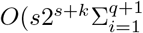 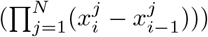 space to find an LCES of *G*_1_, *G*_2_, ..., *G*_*N*_ indexed by *X*_*j*_, where 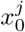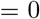 and 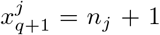 for 1 ≤ *j* ≤ *N*.

## 5 Experiments and analysis

An entire euchromatic human genome is determined by 24 chromosomes, including two sex chromosomes signed as *X* and *Y* and 22 autosomes signed as 1 ~ 22 [25] [26]. A gorilla genome is determined by 25 chromosomes, including the sex chromosomes *X* and *Y* and the autosomes 1, 2*A*, 2*B* and 3 ~ 22. The following understandings are assumed affirmative with respect to human and gorilla genomes. There is no gorilla homologue of the human *Y* -chromosome. The human chromosome 2 tends to be homologous with a fusion of the gorilla chromosomes 2*A* and 2*B* [27]. There is a unique gorilla homologue of a human chromosome except the chromosomes *Y* and 2. Usually, the gorilla homologue of a human chromosome refers to the gorilla chromosome in the same sign as the human chromosome.

In expectation of finding order conserved sequences of significant subsequences in human (resp. gorilla) genome, we performed experiments where Index-LCES(*A*, *B*, *X*, *Y*) was employed to quest for solutions aiming at human chromosomes and their gorilla homologues.

### 5.1 Longest order conserved gene subsequences

Based on Index-LCES(*A*, *B*, *X*, *Y*), we developed a Java software that has been available for uses on a Windows (64-bit) desktop PC with an Intel(R) Core(TM) 3 GHz CPU and 16 GB memory. We downloaded from Ensembl genome browser (http://asia.ensembl.org/index.html) the annotation files of ”hg38” and ”gorGor4” by which we constituted a human as well as a gorilla genome composed of gene sequences. By the annotations of ”hg38”, those genes in a human chromosome can be picked out from the chromosome and lined up into a sequence in the same order as they occur in the chromosome. Those genes in a gorilla chromosome can be constituted into a sequence by the annotations of ”gorGor4” in the same way. We refer to a gene sequence constituted in this way from a human (resp. gorilla) chromosome as the *summary* of the chromosome. Any gene in a genome can be represented by a symbol that depends on its name in the annotation file of the genome. Any respective summaries of two given chromosomes can be treated as two linear genomes for LCES(*A*, *B*, *X*, *Y*) to quest for their LCESs indexed by some exemplar subsequences. Two genes are accepted homologous if they are named the same in the annotation files. Thus every time we mention chromosome summaries in what follows, they indicate linear genomes or gene sequences in which two genes are identical and represented by the same symbol if they are named the same in the annotation files of the genomes including them.

Since the human *Y* -chromosome was believed to reject any gorilla homologue [27], we gave up to quest for an LCES of the human *Y* -chromosome and any other one. For simplicity, we would like to treat the fusion of the gorilla chromosomes 2*A* and 2*B* as *the gorilla chromosome* 2 and will mention by *the gorilla homologue* of the human chromosome 2 to indicate it. Thus we face with 23 human/gorilla chromosome pairs whose summary pairs have been given in S1 File. The lengths of these 23 human/gorila chromosome summary pairs are given in Table 1.

**Table 1.** Lengths of human/gorilla chromosome summaries

|  | chr1 | chr2 | chr3 | chr4 | chr5 | chr6 | chr7 | chr8 | chr9 | chr10 | chr11 | chr12 |
| --- | --- | --- | --- | --- | --- | --- | --- | --- | --- | --- | --- | --- |
| human | 5475 | 4200 | 3188 | 2657 | 2988 | 3064 | 3014 | 2485 | 2333 | 2336 | 3364 | 3055 |
| gorilla | 2947 | 2022 | 1716 | 1223 | 2094 | 1542 | 1440 | 1107 | 1165 | 1127 | 1728 | 1506 |
|  | chr13 | chr14 | chr15 | chr16 | chr17 | chr18 | chr19 | chr20 | chr21 | chr22 | chrX |  |
| human | 1402 | 2287 | 2219 | 2558 | 3059 | 1244 | 2991 | 1458 | 875 | 1388 | 2425 |  |
| gorilla | 611 | 1102 | 991 | 1144 | 820 | 474 | 1668 | 787 | 307 | 645 | 1315 |  |

A "huamn" or "gorilla" statistic represents the gene number of a human or gorilla chromosome summary.

Those 23 human/gorilla chromosome summary pairs were all verified solvable for LCES(*A*, *B*, *X*, *Y*) with *X* = *Y* = ”” to reach their LCESs. These LCESs have been prepared ready in S1 File. Since the length of an LCES of two chromosome summaries reflects the structure similarity of them, we present in Table 2 the first row the lengths of those 23 LCESs. Moreover, we present in Table 2 the second as well as the third row all length ratios of those 23 LCESs to the human/gorilla chromosome summaries pair by pair. The running time statistics for the algorithm to take in questing for solutions are presented in Table 2 the fourth row.

**Table 2.** Characteristics of 23 longest common exemplar subsequences

|  | chr1 | chr2 | chr3 | chr4 | chr5 | chr6 | chr7 | chr8 | chr9 | chr10 | chr11 | chr12 |
| --- | --- | --- | --- | --- | --- | --- | --- | --- | --- | --- | --- | --- |
| length | 1658 | 657 | 954 | 598 | 467 | 855 | 599 | 391 | 476 | 431 | 1063 | 542 |
| lces/lh | 0.303 | 0.156 | 0.299 | 0.225 | 0.156 | 0.279 | 0.199 | 0.157 | 0.204 | 0.185 | 0.316 | 0.177 |
| lces/lg | 0.563 | 0.325 | 0.556 | 0.489 | 0.223 | 0.554 | 0.416 | 0.353 | 0.409 | 0.382 | 0.615 | 0.360 |
| time(ms) | 3237 | 669 | 833 | 359 | 243 | 647 | 483 | 308 | 335 | 323 | 1098 | 569 |
|  | chr13 | chr14 | chr15 | chr16 | chr17 | chr18 | chr19 | chr20 | chr21 | chr22 | chrX |  |
| length | 281 | 428 | 470 | 664 | 231 | 179 | 1114 | 463 | 160 | 358 | 652 |  |
| lces/lh | 0.200 | 0.187 | 0.212 | 0.260 | 0.076 | 0.144 | 0.372 | 0.318 | 0.183 | 0.258 | 0.269 |  |
| lces/lg | 0.460 | 0.388 | 0.474 | 0.580 | 0.282 | 0.378 | 0.668 | 0.588 | 0.521 | 0.555 | 0.496 |  |
| time(ms) | 121 | 324 | 213 | 430 | 100 | 95 | 1173 | 241 | 62 | 161 | 387 |  |

A "length" statistic represents the gene number of an LCES. An "lces/lh" (resp. "lces/lg") statistic represents the ratio of the gen number of an LCES to the gene number of a human (resp. gorilla) chromosome summary. A "time(ms)" statistic represents the real running time.

There exist 1 658 distinct named genes in the LCES for the algorithm to reach in questing the summaries of human chromosome 1 and its gorilla homologue. It took 3.237 minutes for the algorithm to reach this LCES, longer than the running time it took to reach any other one. The 23 LCESs have 13 691 genes in total, about 0.228 times as many genes as those 23 human chromosome summaries have (60 065 genes) and 0.464 times as many genes as those 23 gorilla chromosome summaries have (29 481 genes).

### 5.2 Longest order conserved subsequences of pseudo genes

To break through the restriction of genes that have been annotated, we turn attention to find order conserved sequences of significant subsequences in human or gorilla chromosomes without any regard to the annotation files. We refer to a consecutive DNA subsequence as a *pseudo gene*, if it has at least a stated number (500) of DNA bases and occurs in a human chromosome as well as its gorilla homologue. We start with a DNA sequence represented human chromosome and its gorilla homologue to pursue one of their pseudo gene sequence represented LCESs.

A modified version of SDquest [23] was employed to extract two respective pseudo gene subsequences from a human chromosome and its gorilla homologue. For SDquest with input to be a human chromosome and its gorilla homologue, it will output two pseudo gene sequence represented subsequences of the human chromosome and its gorilla homologue respectively. SDquest was set to identify a consecutive DNA subsequence in a human (resp. gorilla) chromosome as a *pseudo gene* if it has at least 500 DNA bases and occurs in the chromosome’s gorilla (resp. human) homologue *with at least* 95% *identity* ^1^ [23]. All pseudo genes for SDquest to identify were encoded with integers and two pseudo genes were treated as *identical* and encoded with the same integer, if they are with at least 95% identity.

The ”hg38” assembly [28] and the ”gorGor4” assembly [29] on UCSC Genome Browser (https://genome.ucsc.edu/) were chosen as human and gorilla genomes. For SDquest to be input with those 23 huamn/gorilla chromosome pairs that cover the human chromosomes 1 ~ 22 and their gorilla homologues as well as the human *X*-chromosome and its gorilla homologue, we were given by SDquest 23 pairs of pseudo gene sequences, that have been prepared ready in S2 File as well as on https://github.com/ShuZhang-sdu/LCES/blob/master/LCESdata.rar. In what follows, we refer to a pseudo gene sequence represented subsequence of a chromosome as a (*pseudo gene*) *summary* of that chromosome. We present in Table 3 the pseudo gene summary lengths of those 23 human chromosomes and their gorilla homologues.

**Table 3.** Lengths of 46 chromosomes

|  | chr1 | chr2 | chr3 | chr4 | chr5 | chr6 | chr7 | chr8 | chr9 | chr10 | chr11 | chr12 |
| --- | --- | --- | --- | --- | --- | --- | --- | --- | --- | --- | --- | --- |
| Human | 7715 | 11718 | 5907 | 6480 | 2661 | 5060 | 5888 | 5425 | 6970 | 5332 | 4141 | 5763 |
| Gorilla | 7296 | 8117 | 6184 | 5944 | 2820 | 5149 | 5199 | 4273 | 4864 | 4313 | 4145 | 4249 |
|  | chr13 | chr14 | chr15 | chr16 | chr17 | chr18 | chr19 | chr20 | chr21 | chr22 | chrX |  |
| Human | 2595 | 3785 | 2981 | 3449 | 473 | 2791 | 1784 | 1672 | 1045 | 1360 | 5352 |  |
| Gorilla | 2708 | 2844 | 2863 | 3082 | 500 | 2293 | 1771 | 1702 | 1053 | 1331 | 5403 |  |

A "Human" or "Gorilla" statistic represents the length of a human or gorilla pseudo gene summary.

#### 5.2.1 LCES without indexed genes

Those 23 pseudo gene summary pairs have been input to LCES(*A*, *B*, *X*, *Y*) with *X* = *Y* = ”” for their LCESs. There are 20 pseudo gene summary pairs whose LCESs were informed to be found by the algorithm. These 20 summary pairs cover those of the human chromosomes 1, 3 ~ 6, 8 ~ 15, 17 ~ 22, *X* and their gorilla homologues whose LCES lengths as well as the running time the algorithm took to reach them, are given in Table 4. We were informed *out of memory* by LCES(*A*, *B*, ””, ””) for it to quest the other three summary pairs.

**Table 4.** The LCES length and the running time to reach the LCES.

|  | chr1 | chr3 | chr4 | chr5 | chr6 | chr8 | chr9 | chr10 | chr11 | chr12 |
| --- | --- | --- | --- | --- | --- | --- | --- | --- | --- | --- |
| length | 6650 | 5742 | 5419 | 2572 | 4790 | 3277 | 3706 | 3003 | 3872 | 3757 |
| time(ms) | 184223 | 3121200 | 69096 | 9086 | 74185 | 18578 | 250506 | 2859650 | 2047407 | 97843 |
|  | chr13 | chr14 | chr15 | chr17 | chr18 | chr19 | chr20 | chr21 | chr22 | chrX |
| length | 2531 | 2418 | 2566 | 457 | 1987 | 1697 | 1615 | 952 | 1147 | 4935 |
| time(ms) | 9150 | 8494 | 10551 | 244 | 5182 | 3499 | 3012 | 890 | 13252 | 53500 |

A "length" statistic represents the length of an LCES. A "time(ms)" statistic represents the real time for the algorithm to get an LCES.

#### 5.2.2 LCES with indexed genes

Prior to invoking Index-LCES(*A*, *B*, *X*, *Y*) with non null *X* and *Y*, we were asked to prepare an exemplar subsequence *X* of *A* and an exemplar subsequence *Y* of *B* where *X* should be identical to *Y*. Let *A* denote a human chromosome pseudo gene summary and *B* the pseudo gene summary of *A*’s gorilla homologoue. Let *R*[*A*] (resp. *R*[*B*]) denote the subsequence of *A*(resp. *B*) that reserves the pseudo genes in *A*(resp. *B*) other than those that occur exactly once in *A* or in *B*. Let *S* be a longest common subsequence of *R*[*A*] and *R*[*B*], that can be found using the usually used textbook algorithm. Then a longest exemplar subsequence of *S* can be extracted by selecting one from those pseudo genes of the same gene family in *S* for all pseudo gene families. A longest exemplar subsequence of a longest common subsequence of *R*[*A*] and *R*[*B*] will be abbreviated as an LESLCS of *A* and *B*. Let *C* be an LESLCS of *A* and *B*. Then an exemplar subsequence of *A*(resp. *B*) identical to *C* can be identified trivially. Let *X*[*C*] (resp. *Y* [*C*]) denote an exemplar subsequence of *A*(resp. *B*) that is identical to *C*. Then there must exist a unique subsequence of *X*[*C*] (resp. *Y* [*C*]) that is identical to a given subsequence of *C*. Let for a subsequence *C*′ of *C*, *X*[*C*][*C*′] (resp. *Y* [*C*][*C*′]) denote the unique subsequence of *X*[*C*] (resp. *Y* [*C*]) that is identical to *C*′. Then *X*[*C*][*C*′] and *Y* [*C*][*C*′] are available for Index-LCES(*A*, *B*, *X*[*C*][*C*′], *Y* [*C*][*C*′]) to quest *A* and *B* for an LCES indexed by *X*[*C*][*C*′] as well as *Y* [*C*][*C*′].

For *C*′ to be a subsequence of *C*, Index-LCES(*A*, *B*, *X*[*C*][*C*′], *Y* [*C*][*C*′]) either runs to reach an LCES of *A* and *B* indexed by *X*[*C*][*C*′] as well as *Y* [*C*][*C*′] or runs into *out of memory*. An execution of Index-LCES(*A*, *B*, *X*[*C*][*C*′], *Y* [*C*][*C*′]) is *successful*, if it happens to reach an LCES of *A* and *B* indexed by *X*[*C*][*C*′] as well as *Y* [*C*][*C*′]. What we wondered first lies on how many pseudo genes a subsequence *C*′ of *C* should have in order for an execution of Index-LCES(*A*, *B*, *X*[*C*][*C*′], *Y* [*C*][*C*′]) to be successful.

What we did for this uncertainty is: For *r* to take value in (0, 1) increasingly, randomly select a subsequence *C*′ of *C* with *r|C|* pseudo genes and examine Index-LCES(*A*, *B*, *X*[*C*][*C*′], *Y* [*C*][*C*′]) for if its execution could be successful. For *r* with 0 *< r <* 1, a subsequence *C*′ of *C* with *C*′ = *r|C|* can be selected randomly for multiple times such that Index-LCES(*A*, *B*, *X*[*C*][*C*′], *Y* [*C*][*C*′]) can be assumed to output a distinct solution each time. In order to catch hold of the real effect of *C*′ on the output of Index-LCES(*A*, *B*, *X*[*C*][*C*′], *Y* [*C*][*C*′]), we performed 10 times as a *round of executions* of Index-LCES(*A*, *B*, *X*[*C*][*C*′], *Y* [*C*][*C*′]) for randomly selected *C*′ in the same length so that the successful executions could be counted up.

We were left with 3 pseudo gene summary pairs that are of the human chromosomes 2, 7 and 16 and their respective gorilla homologues for which we met executions of LCES(*A*, *B*, ””, ””) that are not successful. Their LESLCSs have been prepared ready and collected in S3 File and have 157, 37 and 94 pseudo genes respectively. For all pair *A* and *B* in these three pseudo gene summary pairs, for all *r* ∈ {1%, 10%, 20%, 30%}, we performed a round of executions of Index-LCES(*A*, *B*, *X*[*C*][*C*′], *Y* [*C*][*C*′]) where each time of the algorithm’s execution in a round was with *C*′ randomly selected from *C* and |*C*′ = *r*|*C*|.

In Table 5, we present the respective successful numbers in all rounds of algorithm executions. Those statistics in Table 5 inform that as *r* or the number of pseudo genes in *C*′ increases, the number of successful executions of Index-LCES(*A*, *B*, *X*[*C*][*C*′], *Y* [*C*][*C*′]) increases until its upper limit 10. This also implies it workable for Index-LCES(*A*, *B*, *X*, *Y*) to be used in finding some common exemplar subsequences of two arbitrary linear genomes.

**Table 5.** Successful numbers in 10 times of algorithm invokes

| $r$ | 1% | 10% | 20% | 30% |
| --- | --- | --- | --- | --- |
| chr2 | 0 | 0 | 0 | 6 |
| chr7 | 0 | 8 | 9 | 10 |
| chr16 | 0 | 8 | 10 | 10 |

An entry indicates the number of successful executions of $\text{Index-LCES}(A, B, X[C][C'], Y[C][C'])$ in 10 times of the algorithm executions where $C'$ is a randomly selected subsequence of $C$ with $r|C|$ pseudo genes for a percentage value $r$ .

One should be worried about whether the algorithm could reach a long enough common exemplar subsequence of a pseudo gene summary pair, because more indexed genes used might cause Index-LCES(*A*, *B*, *X*[*C*][*C*′], *Y* [*C*][*C*′]) to reach a solution with less pseudo genes. If an LCES of a pseudo gene summary pair *A* and *B* can be reached by LCES(*A*, *B*, ””, ””), this can be assessed by examining the length ratio of a solution for Index-LCES(*A*, *B*, *X*[*C*][*C*′], *Y* [*C*][*C*′]) to reach to an LCES for LCES(*A*, *B*,””, ””) to reach. Otherwise, since an LCES of two arbitrary pseudo gene summaries cannot be longer than their longest common subsequence, we adopted to assess the length of an algorithm solution by examining the length ratio of a solution for Index-LCES(*A*, *B*, *X*[*C*][*C*′], *Y* [*C*][*C*′]) to reach to a longest common subsequence of *A* and *B*.

We selected three human chromosomes that are signed as 1, 2 and *X* and their gorilla homologues and performed the following examinations: For every pair *A* and *B* of the pseudo gene summary pairs of these three human/gorilla chromosome pairs, for *r ∈* {30%, 35%, 40%, 45%, 50%}, Index-LCES(*A*, *B*, *X*[*C*][*C*′], *Y* [*C*][*C*′]) was invoked repeatedly until the successful algorithm executions came to 10 times, where each algorithm execution was with *C*′ randomly selected from *C* and |*C*′| = *r*|*C*|.

We present in Table 6 the 2^*nd*^, 4^*th*^ and 6^*th*^ columns the average lengths of the solutions for Index-LCES(*A*, *B*, *X*[*C*][*C*′], *Y* [*C*][*C*′]) to reach in questing the three pseudo gene summary pairs. We present in Table 6 the 3^*rd*^ and 5^*th*^ columns the average length ratios of the solutions Index-LCES(*A*, *B*, *X*[*C*][*C*′], *Y* [*C*][*C*′]) had reached in questing the pseudo gene summary pairs of the human chromosomes 1 and *X* and their gorilla homologues to the LCESs of the respective pseudo gene summary pairs.

**Table 6.**
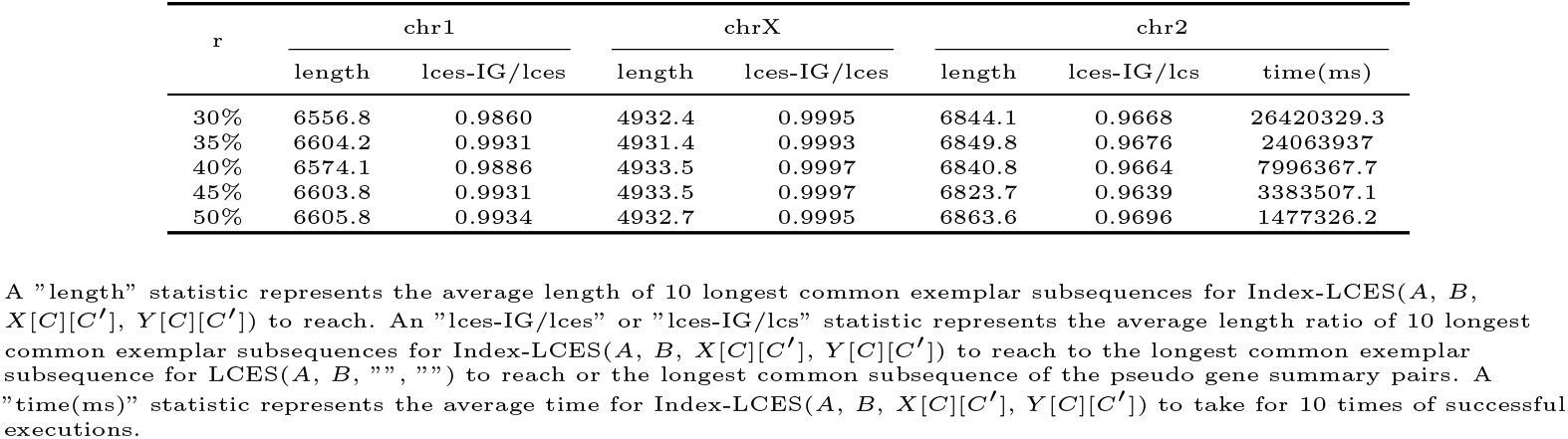
Length ratios of longest common exemplar subsequences

Moreover, we present in Table 6 the 7^*th*^ column the average length ratios of solutions Index-LCES(*A*, *B*, *X*[*C*][*C*′], *Y* [*C*][*C*′]) had reached in questing the pseudo gene summary pair of the human chromosome 2 and their gorilla homologue to the longest common subsequence (7079 pseudo genes) of this two pseudo gene summaries. The statistics in Table 6 show that although there are an increasing number of indexed genes to be selected, there exists no downward trend on the lengths of solutions for the algorithm to reach. The lengths of solutions the algorithm had reached are always very close to the length of a real LCES. This implies that one is allowed to select more indexed genes to make the algorithm take less time to reach a common exemplar subsequence whose length can come very close to an LCES.

As *r* takes value from 30% to 50%, the average running time for Index-LCES(*A*, *B*, *X*[*C*][*C*′], *Y* [*C*][*C*′]) to take in questing the pseudo gene summary pair of the human chromosome 2 and its gorilla homologue, goes down from 440.3 minutes to 24.6 minutes (Table 6 the 8^*th*^ column). In terms of the running speed, the algorithm performed worst in its execution in questing this pseudo gene summary pair for solutions. This might be because there are 12 identical pseudo gene pairs in both of these two pseudo gene summaries whose spans are no less than 4730.

For the pseudo gene summary pairs of the human chromosomes 7 and 16 and their gorilla homologues and for *r* ∈ {10%, 15%, 20%, 25%, 30%}, we present in Table 7 the statistics on how many executions of Index-LCES(*A*, *B*, *X*[*C*][*C*′], *Y* [*C*][*C*′]) we had used in order for the algorithm to run into 10 times of successful executions, how many pseudo genes there are averagely in a solution for a successful execution of the algorithm to reach and how much time a successful execution of the algorithm had taken averagely. Those statistics in Table 7 show that the running time the algorithm had taken averagely to reach a solution rapidly decreases as the value of *r* increases, instead of which, the average length of those common exemplar subsequences for the algorithm to reach remains unchanged.

**Table 7.** LCES of two pseudo gene summary pairs.

|  |  | 10% | 15% | 20% | 25% | 30% |
| --- | --- | --- | --- | --- | --- | --- |
| chr7 | total | 16 | 10 | 10 | 10 | 10 |
|  | length | 4160.8 | 4205 | 4239.7 | 4240.1 | 4239.9 |
|  | time(ms) | 18344280.6 | 4621480.1 | 1415950.3 | 1329840.5 | 322427 |
| chr16 | total | 14 | 14 | 11 | 11 | 10 |
|  | length | 2383.7 | 2376.4 | 2373.9 | 2410.4 | 2418.4 |
|  | time(ms) | 427906.3 | 168763.2 | 133125.5 | 106782.8 | 72637 |

A "total" statistic indicates the total number of $\text{Index-LCES}(A, B, X[C][C'], Y[C][C'])$ executions in which 10 executions were successful. A "length" statistic indicates the average length of those 10 solutions for the 10 successful executions of $\text{Index-LCES}(A, B, X[C][C'], Y[C][C'])$ to reach. A "time(ms)" statistic indicates the average time for the 10 successful executions of $\text{Index-LCES}(A, B, X[C][C'], Y[C][C'])$ to take.

#### 5.2.3 Annotations on pseudo genes in the LCESs

We have collected into S4 File those 20 pseudo gene subsequences for LCES(*A*, *B*, ””, ””) to reach in questing the pseudo gene summary pairs of the human chromosomes 1, 3 ~ 6, 8 ~ 15, 17 ~ 22, *X* and their gorilla homologues.

The pseudo gene summary pairs that are of the human chromosomes 2, 7 and 16 and their gorilla homologues, were verified reachable by Index-LCES(*A*, *B*, *X*, *Y*) with non null *X* as well as *Y*. The LESLCSs of these three summary paires have been stored in S3 File. Three respective subsequences with 48, 4 and 10 pseudo genes of the LESLCSs in S3 File have been stored in S5 File. These three subsequences were obtained by randomly selecting from the three LESLCSs as many pseudo genes as 30%, 10% and 10% the numbers of pseudo genes in them. There are six subsequences of the three pseudo gene summary pairs that fall into three pairs each of which has both of its members identical to one of the three pseudo gene subsequences in S5 File. These six subsequences have also been stored in S5 File. Let *A* and *B* denote an arbitrary pseudo gene summary pair of the human chromosomes 2, 7 and 16 and their gorilla homologues, *C* the LESLCS of *A* and *B* in S3 File and *C*′ the subsequence of *C* in S5 File, *X*[*C*][*C*′] (resp. *Y* [*C*][*C*′]) a subsequence of *A*(resp. *B*) in S5 File that is identical to *C*′. Then *A*, *B*, *X*[*C*][*C*′], *Y* [*C*][*C*′] are available for Index-LCES(*A*, *B*, *X*[*C*][*C*′], *Y* [*C*][*C*′]) to reach an LCES of *A* and *B* indexed by *X*[*C*][*C*′] as well as *Y* [*C*][*C*′]. We were given by Index-LCES(*A*, *B*, *X*, *Y*) three common exemplar subsequences of the pseudo gene summary pairs of the human chromosomes 2, 7 and 16 and their gorilla homologues, which have been collected together into S4 File following the first 20 ones.

An arbitrary pseudo gene in an arbitrary sequence in S4 File can be thought of as conserved as well as significant. We performed experiments where we contrasted the pseudo gene subsequences in S4 File with the annotated genes in human as well as gorilla chromosomes by examining a pseudo gene for if it overlaps with annotated genes. In S6 File (resp. S7 File), we have prepared ready the annotated genes in those 23 human (resp. 24 gorilla) chromosomes that overlap with some pseudo genes in sequences in S4 File.

A pseudo gene in a sequence in S4 File, if does not overlap with any annotated gene, is valuable for people to illuminate its bio-function or significant for people to mine motifs in it. So In S8 File (resp. S9 File), we have prepared ready those pseudo genes that do not overlap with any annotated gene in human (resp. gorilla) chromosomes.

One may be more interested in a pseudo gene, if it both does not overlap with any annotated gene and is long enough. So in S10 File (resp. S11 File), we have prepared ready those pseudo genes that do not overlap with any annotated gene in human as well as gorilla chromosomes and each of them has at least 10 000 bases.

We will mention by CES-2, CES-7 and CES-16 to indicate those three common exemplar subsequences in S4 File that are of the pseudo gene summary pairs of the human chromosomes 2, 7 and 16 and their gorilla homologues. They are of particular concern because they were reached by Index-LCES(*A*, *B*, *X*, *Y*) with some pseudo genes fixed as indexed. Since the gorilla chromosome 2 we mentioned is a fusion of two real gorilla chromosomes, we will give up to show more statistics about CES-2 in contrast with the annotated gene sequences that were extracted from the human chromosome 2 and its gorilla homologue. In Table 8, we disclose more statistic information about CES-7 and CES-16.

**Table 8.**
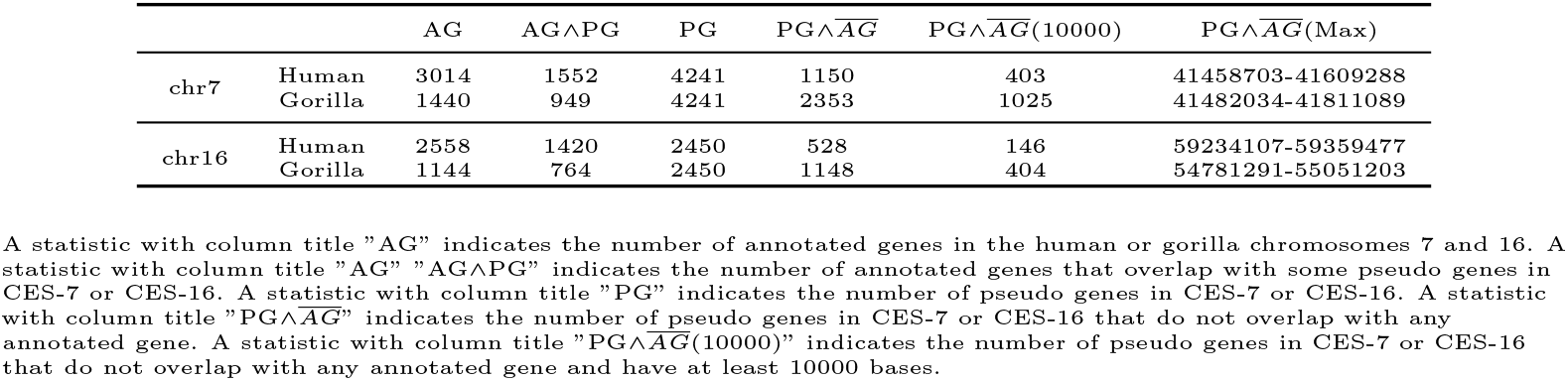
Common exemplar subsequence in contrast with annotated genes

| | | AG | AG$\wedge$PG | PG | PG$\wedge$ $\overline{AG}$ | PG$\wedge$ $\overline{AG}$(10000) | PG$\wedge$ $\overline{AG}$(Max) |
| --- | --- | --- | --- | --- | --- | --- | --- |
| chr7 | Human | 3014 | 1552 | 4241 | 1150 | 403 | 41458703-41609288 |
|  | Gorilla | 1440 | 949 | 4241 | 2353 | 1025 | 41482034-41811089 |
| chr16 | Human | 2558 | 1420 | 2450 | 528 | 146 | 59234107-59359477 |
|  | Gorilla | 1144 | 764 | 2450 | 1148 | 404 | 54781291-55051203 |

A statistic with column title "AG" indicates the number of annotated genes in the human or gorilla chromosomes 7 and 16. A statistic with column title "AG $\wedge$ PG" indicates the number of annotated genes that overlap with some pseudo genes in CES-7 or CES-16. A statistic with column title "PG" indicates the number of pseudo genes in CES-7 or CES-16. A statistic with column title "PG $\wedge$ $\overline{AG}$ " indicates the number of pseudo genes in CES-7 or CES-16 that do not overlap with any annotated gene. A statistic with column title "PG $\wedge$ $\overline{AG}$ (10000)" indicates the number of pseudo genes in CES-7 or CES-16 that do not overlap with any annotated gene and have at least 10000 bases.

In Table 8 the column with title AG (resp. PG), we present the numbers of annotated (resp. pseudo) genes that were extracted from the human chromosomes 7 and 16 and their gorilla homologues. Then in the column with title AG∧PG, we present the numbers of annotated genes in the human chromosomes 7 and 16 and their gorilla homologues, which overlap with pseudo genes in the common exemplar subsequences CES-7 and CES-16. There are 4 241 pseudo genes with 55 185 439 bases in CES-7. Although human genome sequence shows 94.8% similarity to gorilla [30], this common exemplar subsequence has about 0.346 times as many bases as the human chromosome 7 has (159 345 973 bases) and 0.347 times as many bases as the gorilla chromosome 7 has (159 110 946 bases). Moreover, There are 2 450 pseudo genes in CES-16, which have 27 691 501 bases in total, about 0.307 times as many bases as the human chromosome 16 has (90 338 345 bases) and 0.340 times as many bases as the gorilla chromosome 16 has (81 384 781 bases).

To identify new genes or motifs, the pseudo genes that do not overlap with any annotated gene should be more attractive for people to pay attention to. In Table 8 the column with title 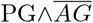, we present the numbers of pseudo genes in CES-7 and CES-16 that do not overlap with any annotated gene extracted from the human chromosome 7 and 16. We present in the column with title 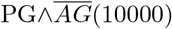 additionally, the numbers of pseudo genes in CES-7 and CES-16 that were involved in the column with title 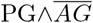 and each of them has at least 10 000 bases. In Table 8 the last column, we present the site intervals of the longest pseudo genes that do not overlap with any annotated gene in CES-7 and CES-16 respectively.

We are informed by these statistics an order conserved sequence of more than 400 (resp. 1 000) pseudo genes all of which are of more than 10 000 bases and do not overlap with any annotated gene in the human (resp. gorilla) chromosome 7. The same two numbers for the human and gorilla chromosome 16 are 140 and 400 respectively.

## Supporting information

### S1 File. The 23 human/gorilla chromosome summary pairs and their longest common exemplar subsequences

This is a compressed file. By uncompressing this file, we can get a folder that contains 23 excel files which are named as ”LCES-chr[*i*].xls” where *i* ∈ {1, ..., 22, *X*}. The excel file with name ”LCES-chr[*i*].xls” gives the summary pair of the human chromosome *i* and its gorilla homologue as well as the LCES of this human/gorilla chromosome summary pair. Each excel file goes along with a table with five columns. Any statistic in the following is assumed in the table whose file name is ”LCES-chr[*i*].xls”. A statistic with column title ”index” indicates the serial number of an annotated gene. A statistic in the column with title ”Human” (resp. ”Gorilla”) indicates the name of a gene in the summary of the human (resp. gorilla) chromosome *i*. An LCES of the summary pair of the human chromosome *i* and its gorilla homologue is concealed in the columns with titles ”Human” and ”signH” (resp. ”Gorilla” and ”signG”). One can verify that the subsequence of annotated genes that occur in the column with title ”Human” accompanied with statistics of ”*” in the column with title ”signH” is identical to the subsequence of genes that occur in the column with title ”Gorilla” accompanied with statistics of ”*” in the column with title ”signG”. Any of these two annotated gene subsequences can be treated as an LCES of the summary pair of the human chromosome *i* and its gorilla homologue.

### S2 File. The pseudo gene summary pairs of 23 human chromosomes and their gorilla homologues

This is a compressed file. By uncompressing this file, we can get a folder that contains 23 text files which are named as ”MosaicSDs SDblockIndexes[*i*].txt” where *i ∈* {1, ..., 22, *X*}. The file with name ”MosaicSDs SDblockIndexes[*i*].txt” gives the pseudo gene summary pair of the human chromosome *i* and its gorilla homologue. Each text file stores pseudo gene sequences in the format of a table with four columns. The statistics in the following are assumed in the file ”MosaicSDs SDblockIndexes[*i*].txt”. A row gives a sequence of multiple pseudo genes and the consecutive DNA subsequence of the human chromosome *i* or its gorilla homologue in which those pseudo genes occur. A statistic in the column with title ”chr” indicates the human or gorilla chromosome in which the data in the same row as it lies in occur. The two statistics in a row with column titles ”start” and ”end” mark the location of the consecutive DNA subsequence in the same row as they lie in in the human or gorilla chromosome *i*. An entry with column title ”SDblocksIndexes” presents all pseudo genes that occur in the consecutive DNA subsequence in the same row as the pseudo genes lie in.

### S3 File. Three LESLCSs of the pseudo gene summary pairs that are of the human chromosomes 2, 7 and 16 and their gorilla homologues

This is a compressed file. By uncompressing this file, we can get a folder that contains three text files which are named as ”LESLCS-chr[*i*].txt” where *i ∈* {2, 7, 16}. The file with name ”LESLCS-chr[*i*].txt” gives the LESLCS of the pseudo gene summary pair of the human chromosome *i* and its gorilla homologue. Each text file contains one line of the pseudo genes separated by commas. The LESLCSs in the aforementioned files are obtained in the following way: Let *A* and *B* denote an arbitrary pseudo gene summary pair of the human chromosomes 2, 7 and 16 and their gorilla homologues, *R*[*A*] (resp. *R*[*B*]) the subsequence of *A*(resp. *B*) that reserves the pseudo genes in *A*(resp. *B*) other than those that occur exactly once in *A* or in *B*. Let *S* be a longest common subsequence of *R*[*A*] and *R*[*B*]. Then a longest exemplar subsequence of *S* is an LESLCS of *A* and *B*.

### S4 File. Common exemplar subsequences of the pseudo gene summary pairs that are of human/gorilla chromosome pairs

This is a compressed file. By uncompressing this file, we can get a folder that contains 23 text files which are named as ”pseudo gene CES-chr[*i*].txt” where *i ∈* {1, ..., 22, *X*}. The file with name ”pseudo gene CES-chr[*i*].txt” gives the common exemplar subsequence of the pseudo gene summary pair that are of the human chromosome *i* and its gorilla homologue. Each text file contains one line of pseudo genes, a common exemplar subsequence.

For a pseudo gene summary pair *A* and *B* of a human chromosome *i ∈* {1, ..., 22, *X*} \ {2, 7, 16} and its gorilla homologue, its LCES is given by LCES(*A*, *B*, ””, ””). For a pseudo gene summary pair *A* and *B* of a human chromosome *i ∈* {2, 7, 16} and its gorilla homologue, for the indexed pseudo gene subsequence pair *X* and *Y* in ”indexed genes-chr[*i*].xls” in S5 File, an LCES of *A* and *B* indexed by *X* as well as *Y* is given by Index-LCES(*A*, *B*, *X*, *Y*).

### S5 File. Three indexed pseudo gene subsequence pairs

This is a compressed file that can be uncompressed into a folder with three excel files which are named as ”indexed genes-chr[*i*].xls” where *i ∈* {2, 7, 16}. The file with name ”indexed genes-chr[*i*].xls” gives the indexed pseudo gene subsequence pair of the pseudo gene summary pair of the human chromosome *i* and its gorilla homologue. Each excel file goes along with a table with three columns whose titles are ”indexedgene”, ”humanpos” and ”gorillapos”. The statistics in the following are assumed in the file ”indexed genes-chr[*i*].xls”. A statistic in the column with title ”indexedgene” gives the identification name of a pseudo gene that is inherited from SDquest and occurs in the pseudo gene summary of the human chromosome *i* as well as the summary of the human chromosome’s gorilla homologue. The statistics in the column with title ”indexedgene” constitute the randomly selected subsequence of the LESLCS in S3 File that is of the pseudo gene summary pair of the human chromosome *i* and its gorilla homologue. Let *A* and *B* denote the pseudo gene summary pair of the human chromosome *i* and its gorilla homologue. A statistic in the column with title ”humanpos” (resp. ”gorillapos”) indicates the identification number in *A*(resp. *B*) of the pseudo gene that is in the same row as it lies in and the column with title ”indexedgene”. Let *X*(resp. *Y*) denote the subsequence of pseudo genes in *A*(resp. whose identification numbers are given in the column with title ”humanpos” (resp. ”gorillapos”). Then the LCES of *A* and *B* indexed by *X* as well as *Y* will be reached by Index-LCES(*A*, *B*, *X*, *Y*) as given in ”pseudo gene CES-chr[*i*].txt” in S4 File.

### S6 File. Annotated genes in 23 human chromosomes that overlap with some pseudo genes

This is a compressed file. By uncompressing this file, we can get a folder that contains 23 excel files which are named as ”AG(∩PG)-human-chr[*i*].xls” where *i* ∈ {1, ..., 22, *X*}. The excel file with name ”AG(∩PG)-human-chr[*i*].xls” gives the annotated genes in the human chromosome *i* that overlap with some pseudo genes in a subsequence that is of the pseudo gene summary of the human chromosome *i* and identical to the common exemplar subsequence in ”pseudo gene CES-chr[*i*].txt” in S4 File. Any statistic in the following is assumed in the table whose file name is ”AG(∩PG)-human-chr[*i*].xls”. A statistic with column title ”index” indicates the serial number of an annotated gene. A statistic in the column with title ”geneID” represents the ID of an annotated gene in the human chromosome *i*. A statistic in the column with title ”genename” represents the name of the annotated gene in the same row as it lies in. The two statistics in a row with column titles ”start” and ”end” mark the location of the annotated gene in the same row as they lie in in the human chromosome *i*. An entry with column title ”pseudogene” presents all pseudo genes that overlap with the annotated gene in the same row as the pseudo genes lie in.

### S7 File. Annotated genes in 24 gorilla chromosomes that overlap with some pseudo genes

This is a compressed file. By uncompressing this file, we can get a folder that contains 24 excel files which are named as ”AG(∩PG)-gorilla-chr[*i*].xls” where *i ∈* {1, 2*A*, 2*B*, 3, ..., 22, *X*}. For *i ∈* {1, 3, ..., 22, *X*}, the file with name ”AG(∩PG)-gorilla-chr[*i*].xls” gives the annotated genes in the gorilla chromosome *i* that overlap with some pseudo genes in a subsequence that is of the pseudo gene summary of the gorilla chromosome *i* and identical to the common exemplar subsequence in ”pseudo gene CES-chr[*i*].txt” in S4 File. For *i* ∈ {2*A*, 2*B*}, the file with name ”AG(∩PG)-gorilla-chr[*i*].xls” gives the annotated genes in the gorilla chromosome *i* that overlap with some pseudo genes in a subsequence that is of the pseudo gene summary extracted from the gorilla chromosome 2 and identical to the common exemplar subsequence in ”pseudo gene CES-chr[2].txt” in S4 File. Each excel file goes along with a table with six columns. Any statistic in the following is assumed in the table whose file name is ”AG(∩PG)-gorilla-chr[*i*].xls”. A statistic with column title ”index” indicates the serial number of an annotated gene. A statistic in the column with title ”geneID” represents the ID of an annotated gene in the gorilla chromosome *i*. A statistic in the column with title ”genename” represents the name of the annotated gene in the same row as it lies in. The two statistics in a row with column titles ”start” and ”end” mark the location of the annotated gene in the same row as they lie in in the gorilla chromosome *i*. An entry with column title ”pseudogene” presents all pseudo genes that overlap with the annotated gene in the same row as the pseudo genes lie in.

### S8 File. Pseudo genes in 23 pseudo gene summaries of human chromosomes that do not overlap with any annotated gene

This is a compressed file. By uncompressing this file, we can get a folder that contains 23 text files which are named as ”PG(-AG)-human-chr[*i*].txt” where *i ∈* {1, ..., 22, *X*}. The file with name ”PG(-AG)-human-chr[*i*].txt” gives the pseudo genes in a subsequence that is of the pseudo gene summary of the human chromosome *i* and identical to the common exemplar subsequence in ”pseudo gene CES-chr[*i*].txt” in S4 File such that all these pseudo genes do not overlap with any annotated gene. Each text file stores pseudo genes in the format of a table with three columns. The statistics in the following are assumed in the file ”PG(-AG)-human-chr[*i*].txt”. A statistic with column title ”index” indicates the serial number of a pseudo gene. The two statistics in a row with column titles ”start” and ”end” mark the location of a pseudo gene in the human chromosome *i*.

### S9 File. Pseudo genes in 23 pseudo gene summaries of gorilla chromosomes that do not overlap with any annotated gene

This is a compressed file. By uncompressing this file, we can get a folder that contains 23 text files which are named as ”PG(-AG)-gorilla-chr[*i*].txt” where *i ∈* {1, ..., 22, *X*}. The file with name ”PG(-AG)-gorilla-chr[*i*].txt” gives the pseudo genes in a subsequence that is of the pseudo gene summary extracted from the gorilla chromosome *i* and identical to the common exemplar subsequence in ”pseudo gene CES-chr[*i*].txt” in S4 File such that, all these pseudo genes do not overlap with any annotated gene. Each text file stores pseudo genes in the format of a table with three columns. The statistics in the following are assumed in the file ”PG(-AG)-gorilla-chr[*i*].txt”. A statistic with column title ”index” indicates the serial number of a pseudo gene. The two statistics in a row with column titles ”start” and ”end” mark the location of a pseudo gene in the gorilla homologue of the human chromosome *i*.

### S10 File. Pseudo genes in 23 pseudo gene summaries of human chromosomes that do not overlap with any annotated gene and have at least 10000 bases

This is a compressed file. By uncompressing this file, we can get a folder that contains 23 text files which are named as ”PG(-AG)(10000)-human-chr[*i*].txt” where *i ∈* {1, ..., 22, *X*}. The file with name ”PG(-AG)(10000)-human-chr[*i*].txt” gives the pseudo genes in a subsequence that is of the pseudo gene summary of the human chromosome *i* and identical to the common exemplar subsequence in ”pseudo gene CES-chr[*i*].txt” in S4 File such that all these pseudo genes do not overlap with any annotated gene and each of them has at least 10 000 bases. Each text file stores pseudo genes in the format of a table with three columns. The statistics in the following are assumed in the file ”PG(-AG)(10000)-human-chr[*i*].txt”. A statistic with column title ”index” indicates the serial number of a pseudo gene. The two statistics in a row with column titles ”start” and ”end” mark the location of a pseudo gene in the human chromosome *i*.

### S11 File. Pseudo genes in 23 pseudo gene summaries of gorilla chromosomes that overlap with no annotated gene and have at least 10000 bases

This is a compressed file. By uncompressing this file, we can get a folder that contains 23 text files which are named as ”PG(-AG)(10000)-gorilla-chr[*i*].txt” where *i* ∈ {1, ..., 22, *X*}. The file with name ”PG(-AG)(10000)-gorilla-chr[*i*].txt” gives the pseudo genes in a subsequence that is of the pseudo gene summary extracted from the gorilla chromosome *i* and identical to the common exemplar subsequence in ”pseudo gene CES-chr[*i*].txt” in S4 File such that, all these pseudo genes do not overlap with any annotated gene and each of them has at least 10 000 bases. Each text file stores pseudo genes in the format of a table with three columns. The statistics in the following are assumed in the file ”PG(-AG)(10000)-gorilla-chr[*i*].txt”. A statistic with column title ”index” indicates the serial number of a pseudo gene. The two statistics in a row with column titles ”start” and ”end” mark the location of a pseudo gene in the gorilla homologue of the human chromosome *i*.

## Acknowledgments

This research is supported by National Natural Science Foundation of China under grant 61732009, 61628207, 61872427, National Natural Science Foundation of Shandong Province under grant ZR201702190130.

## Footnotes

1 95% identity means 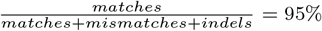, where *matches* (resp. *mismatches*) is the number of matched (resp. mismatched) nucleotides and *indels* is the number of indels.

